# Genome-wide mapping of persistent long-range correlation in the complete telomere-to-telomere human reference genome

**DOI:** 10.64898/2026.07.19.739401

**Authors:** Kleio Papatheodorou, Georgios Megalovasilis, Apostolos Zaravinos, Ilias Georgakopoulos-Soares

## Abstract

Long-range correlations (LRCs) in DNA sequences have been reported for decades, but their interpretation has been limited by incomplete representation of repetitive and structurally complex regions in earlier human genome assemblies. Here, we use the complete telomere-to-telomere (T2T) human reference genome to systematically map LRC structure across all human chromosomes. Most chromosomes exhibited persistent long-range dependence (LRD), with chromosome-specific scaling intervals spanning kilobase to megabase scales and wavelet Hurst exponents consistently above the uncorrelated expectation of 0.5. Spatially resolved Hurst landscapes revealed that this signal is not uniformly distributed along chromosomes, but instead reflects a heterogeneous genomic mosaic. For the purine-pyrimidine encoding, chromosomes 9, 15, X, and Y showed complex fluctuation profiles not adequately summarized by a single scaling exponent; multifractal detrended fluctuation analysis (MF-DFA) confirmed broad, q-dependent multiscaling consistent with multifractal behavior in these chromosomes, consistent with heterogeneous contributions from satellite-rich and structurally distinct sequence compartments. Surrogate analyses further showed that the observed correlations exceed expectations from base composition alone and are not fully explained by local sequence structure preserved in block-shuffled controls. However, short-memory Autoregressive Moving-Average (ARMA) surrogates reproduced the observed H range in a subset of chromosomes, indicating that the strength of evidence for LRD is chromosome-dependent. Together, these results demonstrate that LRC is a widespread but spatially heterogeneous property of the complete human genome. The T2T assembly reveals that previously unresolved repetitive and satellite-rich regions are strongly associated with chromosome-scale and local scaling variation, providing a framework for future studies of genome evolution, chromatin structure, recombination, structural variation, and genomic instability.

## Introduction

The human genome is organized across a broad hierarchy of sequence scales, from local motifs and nucleosome-scale periodicities to megabase-sized compositional domains, segmental duplications, satellite arrays, and chromosome-arm structures. This multiscale organization raises a fundamental question: to what extent does the linear DNA sequence exhibit statistical dependencies that persist across long genomic distances? In this context, fractal or scale-invariant organization refers to signals whose statistical properties remain similar across multiple scales, such that local and global patterns are not independent but are linked through long-range dependencies (LRDs).

Previous work has showcased the existence of long-range correlations (LRCs) in the human genome (Buldyrev et al. 1995; Arneodo et al. 1995; Audit et al. 2001; Peng et al. 1992). Methodologies used to investigate long-distance correlations in DNA sequences have included DNA-walk representations and fluctuation analysis (Peng et al. 1992), power-spectral or 1/f-noise analysis (Li and Kaneko 1992; Voss 1992), mutual information and other information-theoretic measures (Li and Kaneko 1992; Grosse et al. 2000; Holste et al. 2003), wavelet-transform approaches, including wavelet analysis of DNA walks (Haimovich et al. 2006; Arneodo et al. 1995), detrended fluctuation analysis (DFA) and local scaling-exponent methods (Peng et al. 1994; Buldyrev et al. 1995; Carpena et al. 2007), and multifractal approaches, including measure-based multifractal analysis, MF-DFA (Kantelhardt et al. 2002; Rosas et al. 2002; Yu et al. 2001).

Biological interpretations of LRCs have included compositional heterogeneity (Larhammar and Chatzidimitriou-Dreismann 1993), patchiness (Karlin and Brendel 1993), and isochore organization, the hierarchical structural organization of chromatin (Audit et al. 2001), mutational mechanisms (Gu and Li 1994; Li et al. 1994), and transposable elements and repeats (Buldyrev et al. 1993; Holste et al. 2003). However, until recently, the most repetitive and structurally complex regions of the human genome were absent or fragmented in reference assemblies, limiting analyses that require uninterrupted chromosome-scale sequence. The telomere-to-telomere (T2T) human reference genome assembly has now made it possible to study complete human chromosomes as uninterrupted sequences, including regions that were previously unresolved because of high repeat content, such as centromeres, pericentromeric satellites, telomeric regions, acrocentric short arms, and large segmental duplications (Nurk et al. 2022).

Here, we leverage the T2T reference human genome assembly to systematically examine long distance correlations across the human chromosomes. Rather than imposing a single genome-wide scaling exponent, we apply a maximum-likelihood detrended fluctuation analysis (ML-DFA) framework to identify chromosome-specific scaling regimes and distinguish linear, segmented, and complex fluctuation profiles. Repeat-exclusion and local repeat-content analyses show that repeats elevate local and chromosome-level Hurst exponents, but persistent correlations after repeat removal indicate that repeats amplify rather than fully explain genome-wide LRCs. Additionally, we generate both chromosome-wide and sub-chromosomal maps of persistent correlation structure, enabling the localization of regions with unusually strong or weak scaling behavior. We validate observed LRCs against surrogate null models that preserve different aspects of sequence composition and local structure, helping to distinguish genuine long-range organization from effects driven by nucleotide composition, short-range dependencies, repeat content or finite-size biases. We conclude that the complete T2T human genome exhibits chromosome-specific, long-range and multifractal correlation structures that are strongly shaped by repetitive, satellite-rich, and compositionally heterogeneous regions previously inaccessible to genome-wide analysis.

## Materials and Methods

### Sequence encoding and DNA walk

Each chromosome was primarily encoded by the purine-pyrimidine rule: *b*_*t*_ =+1ifnucleotide *s*_*t*_ ∈ {*A, G*}, -1 if *s*_*t*_ ∈ {*C, T*}; positions with ambiguity codes were excluded. For completeness, selected analyses were also repeated using the weak/strong hydrogen-bond encoding, ({A,T}) versus ({G,C}), and the amino/keto encoding, ({A,C}) versus ({G,T}), as complementary sensitivity analyses. The mean centered DNA walk,

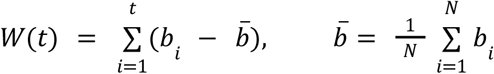

accumulates purine/pyrimidine deviations from global composition. Mean-centering removes the deterministic linear drift induced by global purine/pyrimidine imbalance, so that excursions in W(t) reflect positional organisation around the chromosome-specific composition rather than the marginal nucleotide-class frequencies alone. The purine/pyrimidine projection is a standard binary representation for LRC analysis of DNA sequences (Peng et al. 1992; Voss 1992) and captures the compositional variation that may be related to isochores, repeats, and chromatin-associated genomic domains.

### Scale segmentation by ML-DFA

Vertebrate chromosomes are compositionally heterogeneous, and isochore boundaries, centromeric repeats, and transposable-element clusters each impose distinct scaling regimes on the DNA walk. Therefore, a single DFA exponent fitted across the full scale range may conflate distinct patterns. To avoid imposing a single global power law *a priori*, the ML-DFA algorithm of Botcharova et al. (2013) (Botcharova et al. 2013) is applied, which selects the shape of the log-log fluctuation profile from a candidate model set using the Akaike Information Criterion (AIC).

#### Fluctuation function

For each window size s the DNA walk W(t) is divided into 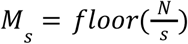 non overlapping windows. Within each window v, a first-order polynomial *p*_*v*_(*i*) was fitted to the cumulative profile and subtracted. The DFA-1 fluctuation function at window size s is computed as,

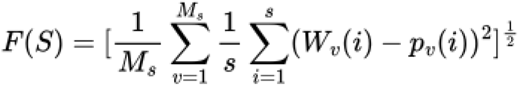

#### Candidate model set

The log-log profile *log*_10_ *F* versus *log*_10_ *s* is min-max normalised to [0,100] and fitted by eight candidate functions: linear, quadratic, cubic, quartic, square-root, logarithmic, two-segment piecewise linear spline and three-segment piecewise linear spline. Polynomial and nonlinear models are fitted by least squares or nonlinear least squares. Piecewise models were fitted by exhaustive grid search over valid breakpoint locations, requiring at least three scale points per segment. Each model was ranked by,

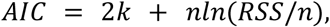

where k is the number of free parameters (k=2,3,4,5,2,2,5,8 for the eight models respectively), n=60 scale points and RSS is the residual sum of squares. The winning model minimises AIC. The linear model was considered statistically indistinguishable from the best model when Δ*AIC* =*AIC*_*linear*_ − *AIC*_*best*_ < 2.

#### Candidate segment extraction

Only winning linear and piecewise-linear models yield candidate scaling segments with interpretable slope estimates. A winning linear model yields one segment spanning the full scale range. Winning two- and three-segment spline models yield two or three candidate segments, respectively, each with its own slope α estimated by linear regression on (*log*_10_ *s, log*_10_ *F*) within the segmented boundaries. Curved models, including quadratic, cubic, quartic, square-root, and logarithmic profiles, are treated as evidence of non-single-regime scaling and do not produce candidate LRD segments. Chromosomes for which ML-DFA selected a quartic or otherwise complex detrending model are not discarded. Instead, these chromosomes are assigned to a separate multifractal analysis branch. This decision reflects the fact that a single Hurst exponent may be insufficient when the fluctuation curve exhibits strong curvature or multiple scaling regimes. For such chromosomes, MF-DFA is used to evaluate whether the observed complexity reflects heterogeneous scaling behavior across small- and large-fluctuation regions.

#### Primary segment selection

Candidate segments are filtered and ranked by the following ordered criteria: i) Structural exclusion: for two- and three-segment spline models, the leftmost segment is excluded as a short-range regime; no positional exclusion is applied for other models; ii) Alpha-range: 0.5 < α < 0.95, retaining persistent LRD regimes while excluding anti-persistent or near-non-stationary behaviour; iii) Minimum log-span: Δ*log*_10_ *s* ≥ 1. 3 decades, ensuring the slope is estimated over a sufficiently broad scale range; iv) RMSR quality gate: root mean square residual of the segment’s linear fit < 0.02 if *log*_10_ *F* units; If no segment satisfied the RMSR threshold but passed the structural, alpha-range, and log-span filters, the segment with the lowest RMSR was retained and flagged for sensitivity analysis; v) Ranking: among qualifying segments, the segment with the highest α is selected as the primary scaling regime, prioritizing the most persistent valid segment. The resulting scale bounds [*s* _*min*_ ^*primary*^, *s* _*max*_ ^*primary*^ ] of the selected segment define the scale interval used for wavelet Hurst estimation and surrogate-based null-model testing.

### Wavelet analysis and spatially resolved Hurst estimation

Wavelet analysis is used as an independent estimator of LRD within the ML-DFA selected scaling interval. For each chromosome, the input signal is the mean-centered purine/pyrimidine binary series 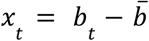 For each retained octave j, the empirical wavelet variance is calculated from the detail coefficients *d*_*j,k*_ as,

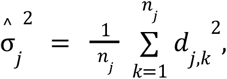

where *n*_*j*_ is the number of coefficients at octave j. For a stationary LRD signal, the wavelet variance scales as,

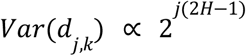

and therefore,

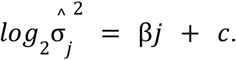

The Hurst exponent is estimated as,

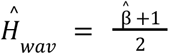

The slope 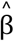 is obtained by weighted least-squares regression of 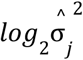 on octave j, using weights proportional to *n*_*j*_. This gives greater weight to finer octaves, where more coefficients are available and variance estimates are more reliable.

The wavelet regression is restricted to the primary scaling interval identified by ML-DFA. The ML-DFA scale bounds [*s* _*min*_ ^*primary*^, *s* _*max*_ ^*primary*^ ] are converted to dyadic octave indices as

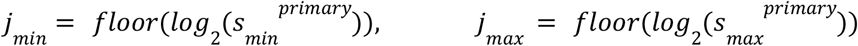

Octaves with fewer than five available detail coefficients are removed to avoid unstable variance estimates. The weighted coefficient of determination R^2^ is recorded as a goodness of fit measure for the log-wavelet variance regression.

To examine spatial heterogeneity in LRD, the wavelet estimator was also applied in overlapping genomic windows. For each chromosome with primary scaling interval [*s* _*min*_ ^*primary*^, *s* _*max*_ ^*primary*^] and corresponding octave range [*j*_*min*_, *j*_*max*_ ], the window length was set to 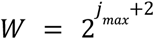. This choice defines a local window large enough to probe the coarsest chromosome-specific octave, while octaves with insufficient coefficient support were excluded within each window using the same filtering criterion as in the chromosome-wide analysis. Windows were advanced in steps of S=W/4 (75% overlap between adjacent windows). At each starting position p, the local mean-centered binary segment was passed to the wavelet estimator using the same octave range [*j*_*min*_, *j*_*max*_ ], as the global estimate, so that the local estimate,

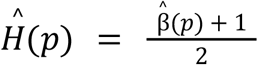

is directly comparable to the chromosome-wide value. The resulting sliding-window H(p) provides a spatially resolved map of correlation strength along each chromosome. Higher values of H(p) indicate regions with stronger persistent scaling structure, whereas values closer to 0.5 indicate weaker or approximately uncorrelated behavior over the analysis scale range.

### Multifractal detrended fluctuation analysis of quartic-scaling chromosomes

Chromosomes for which ML-DFA selected a quartic detrending model were analysed separately using MF-DFA. For each encoded chromosome-level signal, the mean-centred cumulative profile was divided into non-overlapping segments across a range of scales, using segmentation from both the beginning and end of the sequence to minimise boundary loss. Within each segment, a fourth-order polynomial trend was fitted and removed, and the local detrended variance was calculated. Generalised fluctuation functions Fq(s) were then estimated over a symmetric range of moments, q=-5,…,5, with positive q values emphasizing regions with large fluctuations and negative q values emphasizing regions with small fluctuations. The generalised Hurst exponent h(q) was obtained from the scaling relation *F* _*q*_ (*s*) ∝ *s* ^*h*(*q*)^, while the mass exponent was calculated as τ(*q*) =*qh*(*q*) − 1. The singularity spectrum *f*(α) was derived through a Legendre transform and its width, Δα =α_*max*_ − α _*min*_, was used as the primary measure of multifractal strength. Multifractal behavior was supported by systematic variation of h(q) with q, nonlinearity of τ(q) and a non-zero multifractal spectrum width.

### Repeat masking robustness analysis

To evaluate the contribution of repetitive sequence to chromosome-level LRC, repeat-annotated bases were removed from each chromosome before re-estimating the wavelet Hurst exponent. Repeat intervals were excluded from the purine/pyrimidine-encoded chromosome sequence, and the remaining non-repeat bases were concatenated to form a repeat-excluded sequence for each chromosome. The Hurst exponent was then re-estimated using the same db8 wavelet estimator and the same chromosome-specific ML-DFA scaling interval used for the original chromosome-level analysis. This analysis was interpreted as a sensitivity test for the contribution of annotated repeats, rather than as a biological reconstruction of a repeat-free chromosome, because repeat removal changes genomic spacing and introduces artificial junctions between formerly separated non-repeat regions.

### Local repeat-content analysis

To assess whether elevated local H values were concentrated in repeat-rich regions, the sliding-window H landscape was intersected with repeat annotations. For each local H window, the repeat fraction was calculated as the proportion of bases overlapping annotated repeats. Windows were then stratified into low-repeat (<25% repeat covered), medium-repeat (25-75%), and high-repeat (>75%) categories. The distribution of local Hurst exponents was compared across these categories to determine whether elevated local scaling was concentrated in repeat-rich regions or also present in repeat-poor sequence and the association between repeat fraction and local H was quantified using Spearman correlation. In addition, repeat annotations were summarized by class within each window to identify whether specific repeat compartments, such as LINEs, SINEs, LTR elements, satellites or simple repeats were associated with elevated local Hurst exponents.

### Statistical validation using surrogate null models

A wavelet estimate H>0.5 is necessary but not sufficient evidence of LRC: in a finite, heterogeneous chromosome an estimated Hurst exponent above 0.5 can also arise from purine/pyrimidine composition, local clustering, short-range dependence, repeat elements, or finite-size and estimator bias. Because LRD is an asymptotic property that cannot be proven for a finite sequence, for each chromosome independently it is tested whether the observed wavelet exponent within the ML-DFA primary scaling segment [*s*_*min*_ ^*primary*^, *s* _*max*_ ^*primary*^ ] exceeds that of biologically plausible non-LRD null models. For each null model m, B=999 surrogates are generated, the wavelet exponent is re-estimated on the same interval and the one-sided empirical p-value is computed as,

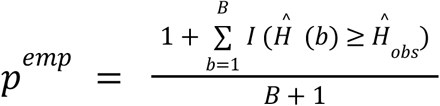

where a small p^emp^ indicates scaling stronger than the null permits and the +1 terms give a valid finite-sample rank-based p-value (Davison and Hinkley 1997).

#### Shuffle surrogates

randomly permute the encoded ± 1series, conserving the purine and pyrimidine counts by destroying all positional structure; they test whether the observed exponent is explainable by composition alone. The null and the alternative hypotheses are,

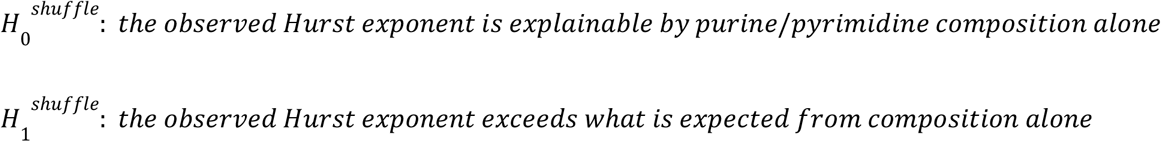

#### Block-shuffle surrogates

were used to test whether the observed scaling could be explained by local sequence structure alone. For each chromosome, the purine/pyrimidine series was partitioned into consecutive non-overlapping blocks of length L, and the order of these blocks was randomly permuted while preserving the internal content of each block. This procedure retains local composition, clustering, and short-range correlations within blocks of size L, but disrupts the ordering of those blocks along the chromosome. Therefore, for each block size L, the null model asks whether the observed wavelet Hurst exponent can be reproduced when genomic structure is preserved only up to the block scale.

Block sizes were chosen from dyadic wavelet scales, L=2^j^, focusing on scales where the sliding-window scalogram indicated spatial heterogeneity, defined by a coefficient of variation CV ≥ 0.05, and requiring 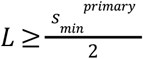. Because correlations within each block are intentionally preserved by the surrogate, the wavelet exponent was re-estimated only on scales larger than the preserved block scale. Specifically, for block size L, the lower bound of the fitting interval was set to,

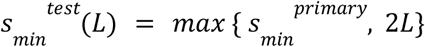

while the upper bound remained,

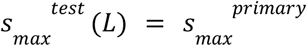

Thus, each block-shuffle test evaluated whether the observed scaling persisted beyond the scale of the preserved blocks, rather than re-measuring structure that the surrogate was designed to retain. Block sizes were retained only when the resulting test interval preserved at least 70% of the original ML-DFA-selected scaling range and produced interpretable wavelet estimates. Fits yielding non-interpretable observed exponents, such as H>1, were excluded from the primary block-size summary. This filtering avoids drawing conclusions from overly narrow or unstable scale intervals, especially for larger block sizes where 2L approaches *s* _*max*_ ^*primary*^.

For each valid chromosome and block size, the observed wavelet Hurst exponent was compared with the empirical distribution of block-shuffle surrogate exponents using a one-sided empirical p-value. Empirical p-values were corrected jointly across all valid chromosome–block-size combinations using the Benjamini–Hochberg false-discovery-rate (BH-FDR) procedure. In addition, for the descriptive summary shown in **Figure 5D**, a chromosome was classified as exceeding the block-shuffle distribution when its observed Hurst exponent was greater than the 97.5th percentile of the corresponding surrogate distribution,

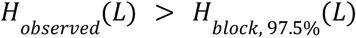

The block-shuffle null hypothesis for block length L is,

*H*_0_ ^*block*^ (*L*) : the observed Hurst exponent is explainable by local structure preserved within blocks of length L. The alternative hypothesis is,

*H*_1_^*block*^ (*L*): the observed Hurst exponent requires ordering beyond block length L.

Persistence of BH-FDR -corrected significance across increasing block sizes was interpreted as evidence that the observed scaling is not explained solely by local composition or short-range clustering. However, results at the largest retained block sizes were interpreted cautiously when only a small number of chromosomes retained valid scale intervals.

For each chromosome and block size, the block-shuffle effect size was calculated as,

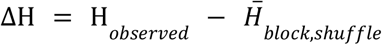

where 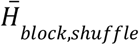 denotes the mean Hurst exponent across the corresponding block-shuffle surrogates. Positive values therefore indicate that the observed chromosome-level scaling exceeded that expected after preserving within-block sequence structure. To evaluate whether ΔH was consistently positive across chromosomes, a one-sided Wilcoxon signed-rank test was additionally performed at each block-size, with Benjamini-Hochberg correction applied across the tested block sizes.

### Autoregressive Moving Average (ARMA) short-memory surrogates

The shuffle and the block-shuffle surrogates address two properties - composition and local clustering. To test whether the observed chromosome-level scaling could be explained by conventional short-memory autocorrelation, we generated B=999 ARMA surrogate sequences. ARMA surrogate analyses were restricted to chromosomes classified as monofractal in the primary scaling analysis. Chromosomes chr9, chr15, chrX, and chrY were excluded from this surrogate framework because their scaling behaviour was better represented by multifractal analyses and was therefore evaluated separately. For each chromosome, the mean-centered purine/pyrimidine series,

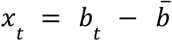

was modelled as an ARMA(p,q) process,

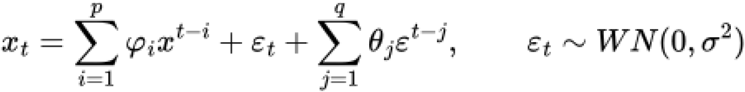

where φ_*i*_ and θ_*j*_ are autoregressive and moving-average parameters and ε_*t*_ is white noise. A stationary ARMA process is a short-memory process whose autocorrelation decays exponentially, in contrast to the power-law decay expected under LRD. Therefore, ARMA surrogates test whether elevated Hurst exponents could arise from finite-order autocorrelation rather than genuine LRD.

The observed exponent was compared with the ARMA surrogate distribution using the one-sided empirical p-value,

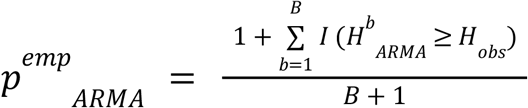

Evidence against the ARMA null was taken when the observed Hurst exponent exceeded the upper tail of the ARMA surrogate distribution.

The null hypothesis is,

H_0_ ^ARMA^: the observed Hurst exponent is consistent with a finite short-memory process with exponentially decaying autocorrelation, whereas the alternative hypothesis is,

H_1_ ^ARMA^: the observed Hurst exponent exceeds what a finite short-memory process produces.

## Results

### ML-DFA analysis across human chromosomes

When a genomic sequence contains multiple compositionally and structurally distinct domains, a single scaling exponent may obscure biologically meaningful heterogeneity in LRC analyses. To determine whether LRD in the complete T2T reference human genome can be described by a single scaling regime or required chromosome-specific segmentation, we first applied ML-DFA independently to each chromosome. This approach allowed us to select the best-fitting fluctuation model for each chromosome and to identify the primary scale interval over which persistent correlation structure was supported. We analyzed three binary nucleotide encodings: purine/pyrimidine, weak/strong hydrogen bonding, and amino/keto. For each chromosome and each encoding, the best-fitting ML-DFA model was selected and the primary scaling interval was defined as the physical scale range over which persistent correlation structure was supported.

Across the three encodings, ML-DFA predominantly selected a three-segment spline model, indicating that chromosome-scale fluctuation profiles generally contained multiple distinct scaling regimes rather than a single linear relationship (**Supplementary Figure 1**). The wavelet Hurst exponent H_wav_ was then estimated from the slope of the wavelet-variance regression over the corresponding admissible wavelet octaves, with the reported R^2^ describing the goodness of fit of this regression. For the purine/pyrimidine encoding, 20 of the 24 chromosomes were best fitted by three-segment splines, whereas chromosomes 9, 15, X, and Y were assigned quartic ML-DFA models (**Table 1; Supplementary Figure 2-3**). Most spline-fitted chromosomes exhibited persistent LRD over chromosome-specific scaling intervals, with Hurst exponent values above 0.5 and strong wavelet-regression fits, whereas chromosomes 9, 15, X, and Y displayed more complex fluctuation profiles and were therefore examined separately using multifractal analysis (**Table 1; Supplementary Figure 4**).

**Table 1:** Summary results across all chromosomes for the purine/pyrimidine encoding. ML-DFA best model selection, scaling interval, DFA scaling exponent α, Wavelet Hurst exponent H and R^2^ goodness of fit are shown. A dash indicates that the corresponding monofractal estimate was not applicable.

| Chromosome | ML-DFA model | Scaling interval | $\alpha$ | $H_{\text{wav}}$ | $R^2$ |
| --- | --- | --- | --- | --- | --- |
| chr1 | Three-segment spline | 370 bp–5.83 Mb | 0.762 | 0.785 | 0.997 |
| chr2 | Three-segment spline | 966 bp–2.17 Mb | 0.728 | 0.782 | 0.992 |
| chr3 | Three-segment spline | 448 bp–84.28 kb | 0.772 | 0.784 | 0.999 |
| chr4 | Three-segment spline | 444 bp–82.34 kb | 0.778 | 0.792 | 0.998 |
| chr5 | Three-segment spline | 346 bp–2.75 Mb | 0.751 | 0.785 | 0.996 |
| chr6 | Three-segment spline | 432 bp–18.67 kb | 0.784 | 0.794 | 0.999 |
| chr7 | Three-segment spline | 679 bp–3.12 Mb | 0.745 | 0.792 | 0.996 |
| chr8 | Three-segment spline | 416 bp–43.58 kb | 0.779 | 0.794 | 0.999 |
| chr9 | Quartic | — | — | — | — |
| chr10 | Three-segment spline | 408 bp–529.10 kb | 0.778 | 0.790 | 0.999 |
| chr11 | Three-segment spline | 408 bp–33.03 kb | 0.776 | 0.790 | 0.999 |
| chr12 | Three-segment spline | 406 bp–26.03 kb | 0.776 | 0.795 | 0.999 |
| chr13 | Three-segment spline | 492 bp–7.72 Mb | 0.815 | 0.815 | 0.996 |
| chr14 | Three-segment spline | 477 bp–171.90 kb | 0.798 | 0.802 | 0.999 |
| chr15 | Quartic | — | — | — | — |
| chr16 | Three-segment spline | 240 bp–3.12 Mb | 0.737 | 0.752 | 0.983 |
| chr17 | Three-segment spline | 292 bp–369.87 kb | 0.752 | 0.787 | 0.990 |
| chr18 | Three-segment spline | 361 bp–60.24 kb | 0.767 | 0.788 | 0.999 |
| chr19 | Three-segment spline | 339 bp–7.17 kb | 0.816 | 0.863 | 0.997 |
| chr20 | Three-segment spline | 534 bp–593.92 kb | 0.736 | 0.768 | 0.992 |
| chr21 | Three-segment spline | 389 bp–121.27 kb | 0.822 | 0.826 | 0.997 |
| chr22 | Three-segment spline | 498 bp–132.98 kb | 0.826 | 0.825 | 0.998 |
| chrX | Quartic | — | — | — | — |
| chrY | Quartic | — | — | — | — |

The weak/strong encoding produced the largest number of complex profiles, with quartic models selected for chromosomes 1, 7, 9, 11, 13, 16, 20, 21, and 22, while the remaining 15 chromosomes were best described by three-segment splines. In contrast, the amino/keto encoding yielded the most consistent segmented behaviour, with three-segment spline models selected for 22 chromosomes and quartic models restricted to chromosomes 17 and 22. This difference suggests that weak/strong nucleotide composition produces more complex chromosome-scale patterns, whereas amino/keto composition shows more consistent scaling behavior. Overall, all three encodings revealed multiple scaling regimes, but the chromosomes showing the most complex fluctuation profiles varied according to the encoding represented.

### Multifractal analysis of Purine/Pyrimidine encoding for chromosomes 9, 15, X, and Y

We next focused our analyses on the purine/pyrimidine encoding because this mapping forms the basis of the original DNA-walk framework for investigating long-range genomic correlations and is a recognized contributor to genomic scaling patterns (Peng et al. 1992; Larhammar and Chatzidimitriou-Dreismann 1993; Buldyrev et al. 1995). In our analysis, it also revealed clear persistent scaling across most chromosomes while identifying a small, well-defined subset with more complex profiles, providing a consistent framework for detailed chromosome-level and multifractal analyses. We observed that chromosomes 9, 15, X, and Y were classified by ML-DFA as exhibiting quartic or otherwise complex scaling behaviour, indicating that their fluctuation profiles are not adequately summarized by a single linear scaling regime (**Supplementary Figure 4**). These chromosomes were therefore analysed separately using MF-DFA (Kantelhardt et al. 2002) to evaluate whether their scale-dependent structure reflected heterogeneous scaling across small- and large-fluctuation regions.

Wavelet scalograms computed across the full available scale range showed increasing variance from fine to coarse scales in all four chromosomes, consistent with persistent scale-dependent structure, with log_2_ wavelet variance increasing monotonically from fine to coarse scales across the entire genomic length (**Figure 1A**). The Hurst exponent H measures the strength of LRD, with H > 0.5 indicating persistent correlations and H = 0.5 indicating an approximately uncorrelated sequence. We report that the global wavelet fits yield H=0.915 (R^2^=0.955), 0.808 (R^2^=0.961), 0.759 (R^2^=0.980), and 0.925 (R^2^=0.908) for chr9, chr15, chrX and chrY respectively, all well above H=0.5, supporting persistent scaling in all four chromosomes. Sliding-window wavelet estimates showed local H(p) values that remained above H=0.5 across the analysed chromosome lengths, generally ranging between approximately 0.55 and 0.70. This supports chromosome-wide persistence over the evaluated scale range, while the stronger local variation observed in chrY is consistent with its more heterogeneous repeat architecture.

**Figure 1:**
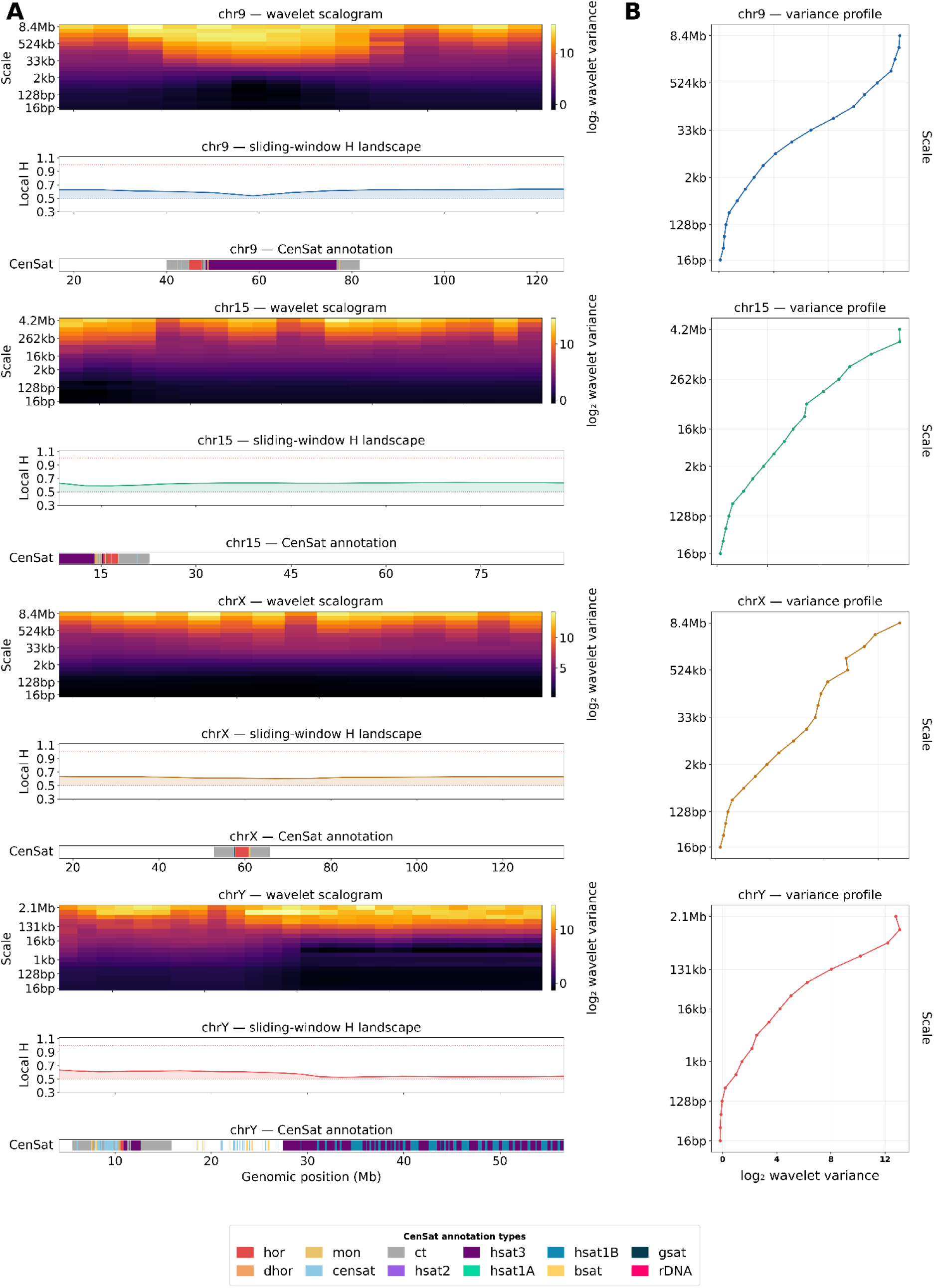
Multiscale wavelet and Hurst analysis across four human chromosomes. **(A)** Wavelet scalograms, sliding-window Hurst landscapes, and CenSat annotation tracks for chr9, chr15, chrX, and chrY; **(B)** Chromosome-level wavelet variance profiles summarize the average octave-wise scaling relationship for each chromosome.

### Satellite-rich regions show suppressed fine-scale wavelet variance

Localized regions of very low fine-scale wavelet variance were visible in chr9 and chrY, primarily over scales from approximately 128 bp to 4 kb. These intervals overlapped satellite-rich regions in the CenSat annotation tracks, including the hsat2/hsat3 block at ∼50-70Mb on chr9 and the hsat3/hsat1B-rich region at ∼26-60 Mb on chrY (**Figure 1**). This pattern is consistent with highly regular tandem repeat structure, whose internal periodicity falls within that scale range, suppressing fine-scale wavelet coefficients. The same transition appears in the MF-DFA fluctuation functions as a sharp discontinuity at *log*_10_(*scale*) ≃ 3 (∼ 1*kb*) **(Figure 2A-D)**, and is most pronounced for the negative-q curves that are sensitive to small fluctuations. These regions do not eliminate local persistence in the wavelet analysis, since local H(p) remains above 0.5, but they likely contribute to the small-fluctuation branch of the multifractal spectrum. Together, these results indicate that satellite-rich tandem repeat arrays define a distinct fine-scale low-variance regime embedded within the chromosome-wide correlation structure, in chromosomes 9 and Y.

**Figure 2:**
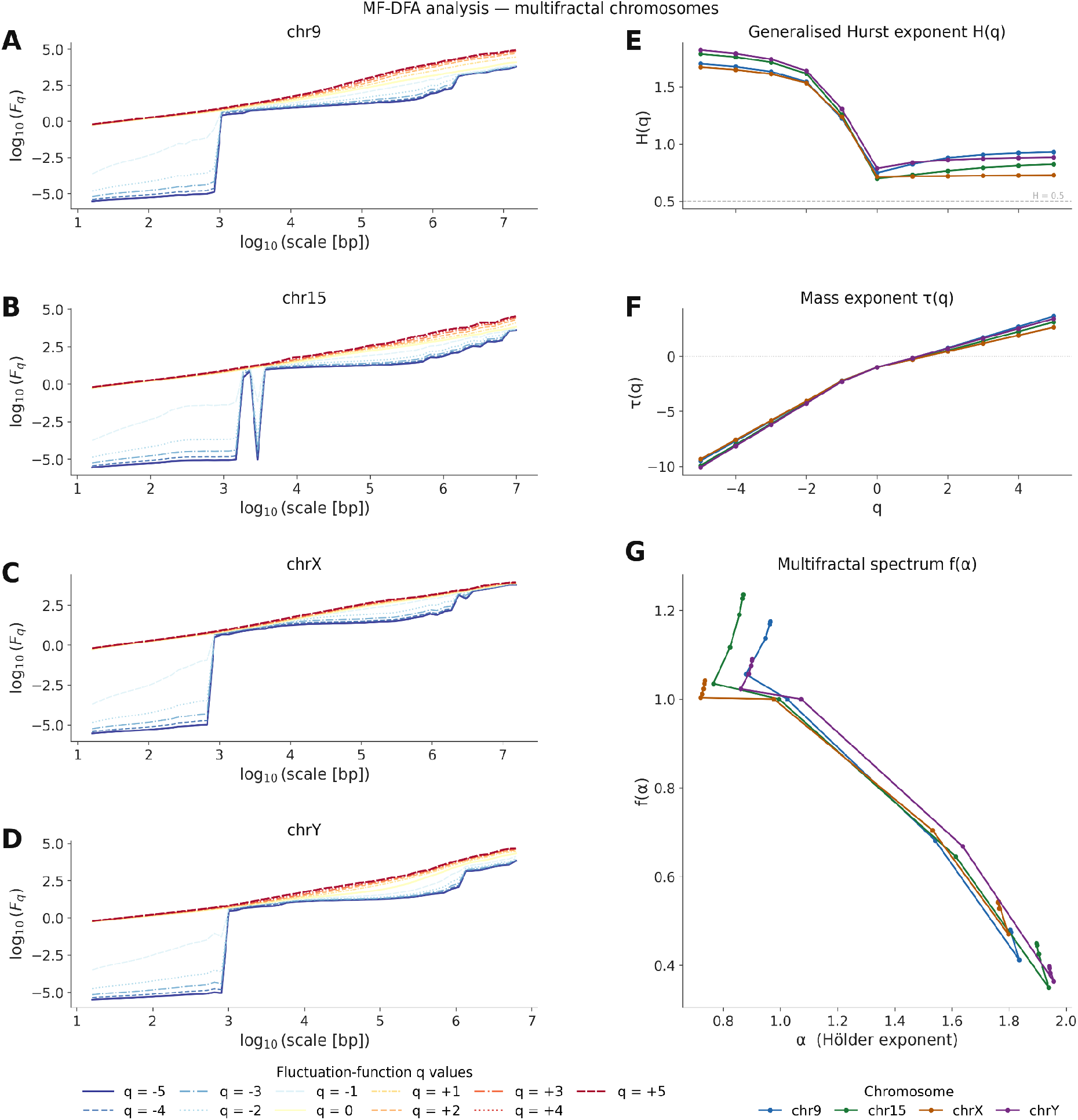
MF-DFA analysis of the four multifractal chromosomes (polynomial order m = 4). **(A-D)** fluctuation functions F*q*(s) for q ∈ [−5, +5]; **(E)** generalised Hurst exponent H(q); (**F)** Mass exponent τ(q); (**G**) Multifractal singularity spectra f(α).

### MF-DFA fluctuation functions reveal heterogeneous scaling

Next, we examined whether different parts of each chromosome scale in the same way across genomic scales. MF-DFA fluctuation functions Fq(s) were computed for *q* ∈ [− 5, 5] using polynomial detrending order (m=4) across all four chromosomes **(Figure 2A-D)**. In all cases the curves fan out into a broad family of clearly distinct slopes, with q=-5 and q=+5 envelopes separated by approximately four orders of magnitude at megabase scales. This fan-out indicates that small- and large-fluctuation genomic regions scale differently with genomic scale, which is the primary empirical signature of multifractal behaviour. A monofractal signal would be expected to produce approximately parallel Fq(s) curves with little dependence of slope on q.

The generalised Hurst exponent H(q), the slope of logFq(s) versus logs, decreases monotonically with increasing q in all four chromosomes **(Figure 2E)**, from *H*(− 5) ≃ 1. 70 − 1. 82 to *H*(+5) ≃ 0. 73 − 0. 93, yielding ΔH = H(-5) - H(+5) of 0.77 (chr9), 0.97 (chr15), 0.95 (chrX) and 0.93 (chrY). This strong q-dependence indicates pronounced heterogeneity in local scaling behaviour. The fact that H(q) remained above 0.5 across the evaluated q-range suggests persistent scaling across both small- and large-fluctuation regimes, although the very high negative-q exponents should be interpreted cautiously because negative moments are sensitive to low-variance regions and finite-size effects. The mass exponent τ(q) is visibly nonlinear in all four chromosomes **(Figure 2F)**, providing additional evidence against a monofractal description independent of H(q): a linear τ(q) would indicate monofractality.

To further characterize the range of local scaling behaviors, we transformed the multifractal exponents into singularity spectra, f(α), which summarize how many different scaling strengths are present in each chromosome. The corresponding singularity spectra f(α) are wide and right skewed (**Figure 2G**) with estimated widths Δα ≃ 0. 96 (chr9), 1.13 (chr15), 1.08 (chrX) and 1.07 (chrY), with most spectra peaking at f(α)=1 near α ≃ 1. 0, consistent with the dominant global wavelet H. The pronounced right skew, the large-α tail being substantially longer than the left arm, reflects the coexistence of qualitatively distinct sequence domains within each chromosome: the gene-rich euchromatic arms, which govern the positive-q scaling and contribute the body of the spectrum near α ≃ 1, and the satellite-rich pericentromeric heterochromatin, which contributes the extreme large-α tail by virtue of its anomalously weak and highly regular local fluctuations. A minor fold at the leftmost spectral points (q < -3) is a recognized numerical artefact of MF-DFA at extreme negative moments and does not affect the interpretation.

Taken together, the four lines of evidence, the fan-out of Fq(s), the strong q-dependence of H(q), the nonlinearity of τ(q), and the broad right-skewed singularity spectra, provide convergent and mutually consistent confirmation that chr9, chr15, chrX, and chrY are not adequately described by a single Hurst exponent. Instead, their scaling behaviour is consistent with multifractal organization, likely reflecting a mixture of genomic domains with different local regularity, including chromosome arms, repeat-rich regions, and satellite-rich pericentromeric intervals.

### Genome-wide spatial heterogeneity of LRC

To determine whether chromosome-level LRC reflects a uniform property of each chromosome or instead emerges from specific genomic compartments, we mapped the local wavelet Hurst exponent across the complete T2T genome and compared this landscape with cytoband, CenSat, RepeatMasker, and GENCODE annotations (**Figure 3**). The genome-wide scalogram (**Figure 3A**) shows scale-dependent variation in wavelet variance across chromosomes, indicating that the strength of sequence organization changes both by genomic position and by scale. Consistent with this, the sliding-window H landscape (**Figure 3B**) shows that persistent scaling is widespread, with most local H values remaining above the uncorrelated reference value of H = 0.5. However, local H is not uniform along chromosomes. Instead, it forms a heterogeneous genomic landscape, with regions of elevated and reduced persistence occurring in distinct chromosomal domains.

**Figure 3:**
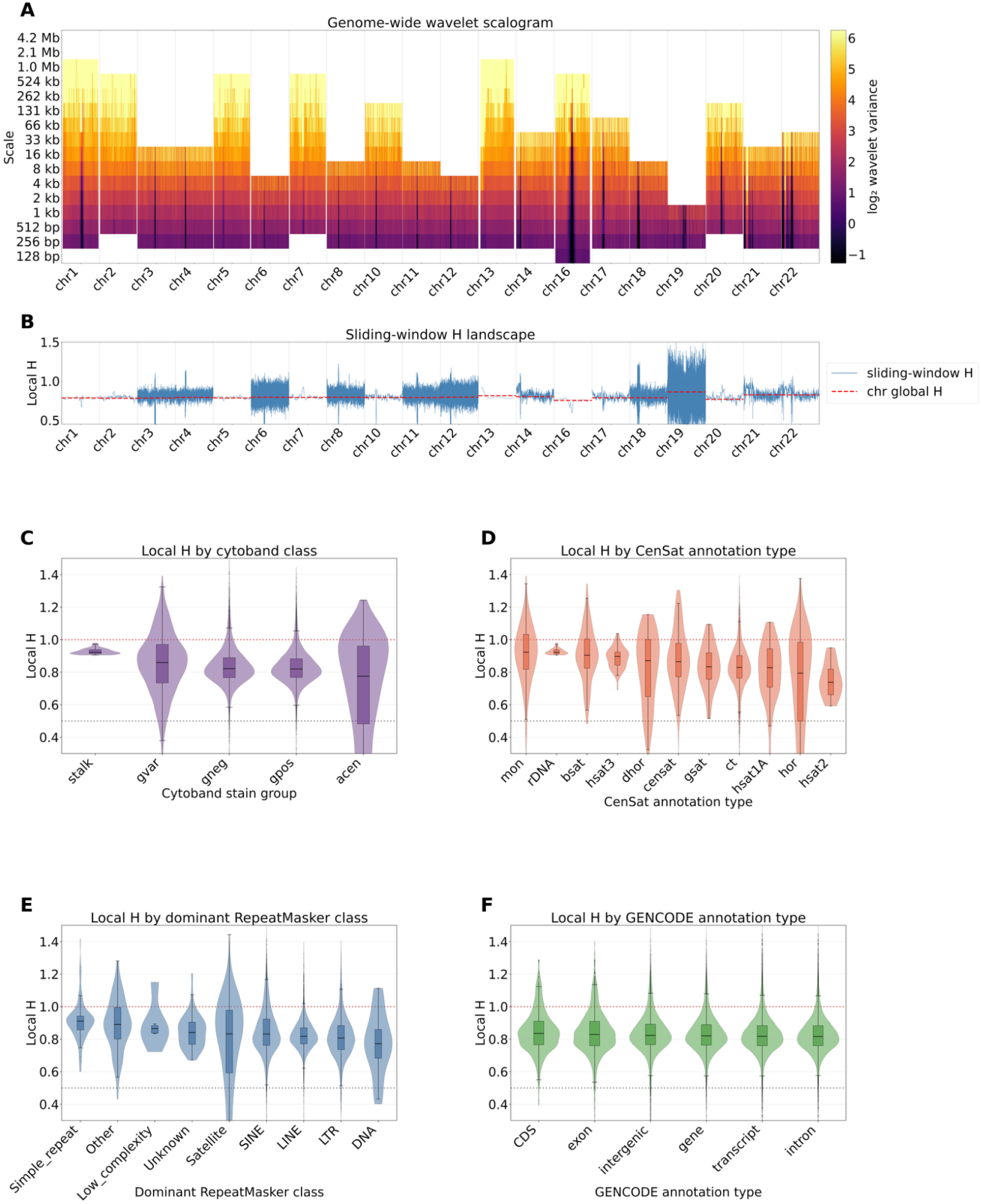
Landscape of local Hurst exponents across genomic sub-compartments and different chromosomes. **(A)** Genome-wide wavelet scalogram, **(B)** Sliding-window local H exponent landscape across genome, **(C)** Distribution of local H values overlapping cytobands, **(D)** Distribution of local H values overlapping CenSat annotations, **(E)** Distribution of local H values overlapping RepeatMasker annotations **(F)** Distribution of local H values across gene annotation features.

This heterogeneity is biologically plausible in the complete T2T assembly, which includes centromeric, pericentromeric, satellite-rich, and repeat-dense regions that were incompletely represented in earlier human reference genomes. Such regions contain long tandem arrays, higher-order repeat structures, and compositionally biased domains that can influence scaling across kilobase-to-megabase ranges. Therefore, repeat-rich compartments are expected to affect local H in a scale-dependent manner: some may increase persistence by introducing extended regularity, whereas highly homogeneous arrays may also produce low-variance regimes at particular scales, as reflected by local transitions in the scalogram.

The annotation-stratified distributions support this interpretation. Cytoband classes differ in their local H distributions (**Figure 3C**), indicating that broad chromosomal compartments are associated with different levels of sequence persistence. CenSat annotations show additional variation among centromere- and satellite-associated sequence classes (**Figure 3D**), suggesting that different repeat architectures contribute differently to the genome-wide correlation landscape. RepeatMasker classes also show distinct local H distributions (**Figure 3E**), further supporting the contribution of repetitive DNA to the observed spatial heterogeneity. In contrast, GENCODE categories show more overlapping distributions across CDS, exons, genes, introns, and transcripts (**Figure 3F**), suggesting that conventional gene-feature classes alone do not explain the strongest variation in local H.

Together, these results support the view that chromosome-level Hurst exponents summarize an underlying heterogeneous genomic landscape rather than a spatially uniform correlation process. While previous studies have linked LRCs in DNA to compositional heterogeneity, repeats, and large-scale genome organization, the complete T2T assembly allows these relationships to be evaluated across previously unresolved satellite-rich and centromeric regions. In this context, the local H landscape suggests that persistent scaling in the human genome reflects the combined contribution of multiple sequence compartments, including satellite-rich centromeric regions, repeat-dense domains, and compositionally structured chromosome arms.

### Repeats contribute only partially to chromosome-level autocorrelations

Because the local H landscape suggested that repeat-rich and satellite-rich compartments contribute strongly to spatial variation in LRC, we next tested how much of the chromosome-level signal remained after excluding annotated repeats. Across all 20 monofractal chromosomes, repeat exclusion reduced the chromosome-level Hurst exponent (**Figure 4A-B**). The median chromosome-level H decreased from 0.791 in the original chromosomes to 0.671 after repeat exclusion, with a median reduction of ΔH=0.121. The reduction was positive for every chromosome, indicating that annotated repeats make a consistent contribution to chromosome-level scaling. However, repeat exclusion did not reduce chromosome-level H to the uncorrelated expectation of 0.5. All repeat-excluded chromosomes retained H values in the range of approximately 0.55-0.70, with chr19 retaining the highest value of H=0.77. This indicates that repeats substantially elevate chromosome-level H, but the persistent scaling signal is not exclusively confined to annotated repeat sequences. Therefore, the repeat-exclusion analysis supports a mixed interpretation: repetitive sequence contributes strongly to LRC, but non-repeat sequence also retains persistent organisation over the analysed scale intervals.

**Figure 4:**
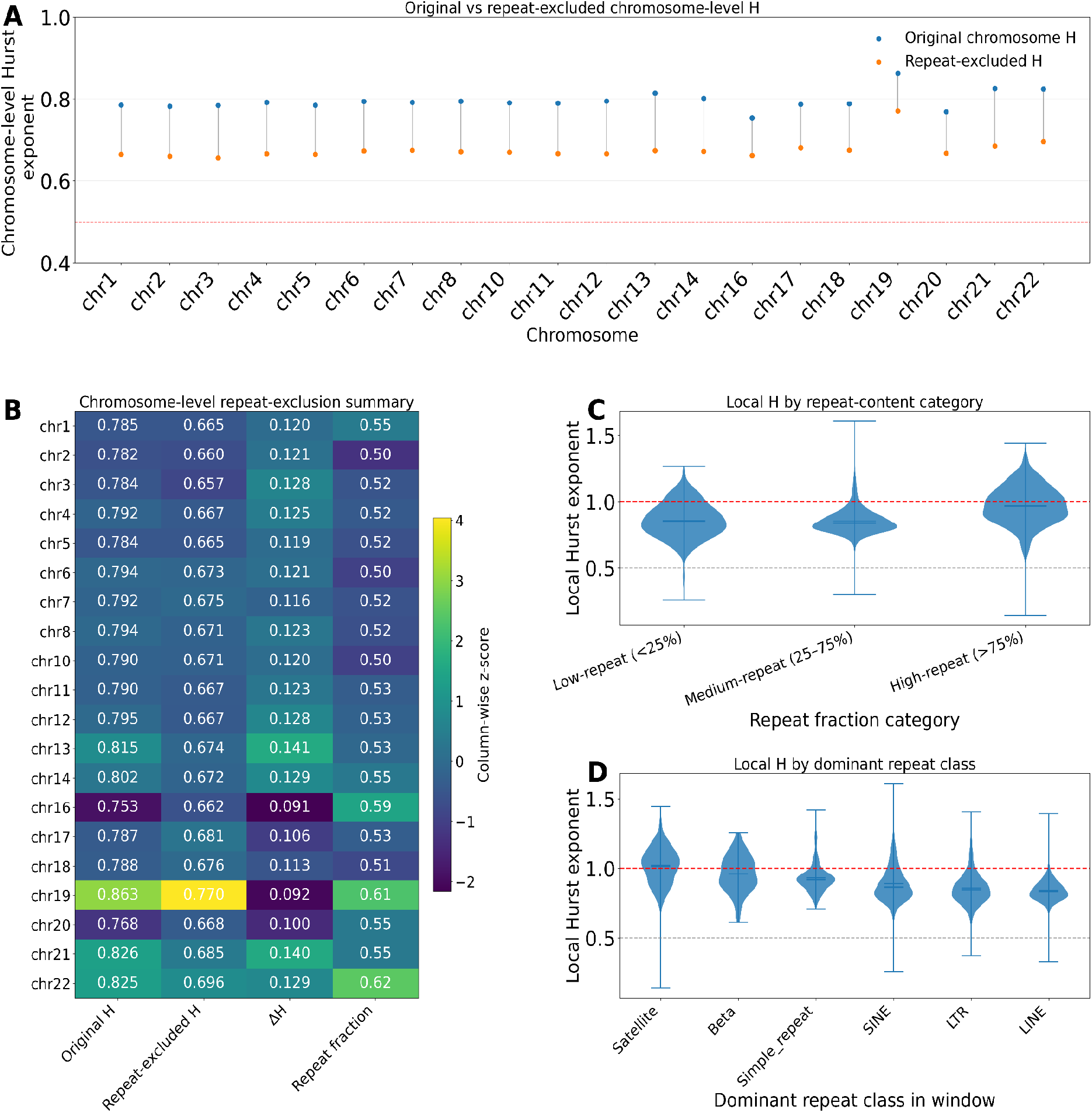
Effects of repeat content on chromosome-level and local Hurst exponents. **(A)** Original and repeat-excluded chromosome-level wavelet Hurst exponents. Blue points show H estimated from the original chromosome sequence, and orange points show H after excluding annotated repeats; **(B)** Chromosome-level repeat-exclusion summary showing original H, repeat-excluded H, ΔH, and chromosome repeat fraction. Numbers printed in the cells are raw values, while colours represent standardized values; **(C)** Distribution of local H values across low-repeat, medium-repeat, and high-repeat windows; **(D)** Distribution of local H values by dominant repeat class within each window.

At the local level, repeat-rich windows showed higher H values than low- and medium-repeat windows (**Figure 4C**). High-repeat windows had the highest median local H, with median H=0.968, compared with 0.852 in low-repeat windows and 0.837 in medium-repeat windows. The association between repeat fraction and local H was positive but moderate, with Spearman ρ=0.279 across 30,548 windows. Dominant repeat-class analysis further shows that the highest local H values were associated with satellite-dominated windows (**Figure 4D**). Satellite windows had the highest median local H, H=1.022, followed by beta-satellite-dominated windows, H=0.96, and simple-repeat-dominated windows, H=0.917. SINE-, LTR-, and LINE-dominated windows had lower median local H values of 0.862, 0.844 and 0.831, respectively. These results indicate that tandem and satellite-rich repeat compartments are particularly associated with elevated local scaling. However, because some repeat classes had a very small number of windows, interpretation should focus primarily on well-represented classes such as Satellite, SINE, LTR and LINE.

Repeating the repeat-content analysis with the complementary encodings revealed a clear encoding dependence (**Supplementary Figure 6-7**). Under amino/keto encoding, repeat exclusion reduced H in all 22 monofractal chromosomes, lowering the median chromosome-level H from 0.857 to 0.712 (median ΔH = 0.145); high-repeat windows and satellite-dominated windows also showed the highest local H values. By contrast, weak/strong encoding showed weak net reduction across its 15 monofractal chromosomes and median H changed from 0.789 to 0.797 (median ΔH = −0.005), with H decreasing in six chromosomes, remaining essentially unchanged in one, and increasing in eight, most strongly in chrY. Local weak/strong H differed only modestly across repeat-content categories and dominant repeat classes.

Together, the repeat-exclusion and local repeat-content analyses show that the contribution of repeats to LRC is encoding-dependent. For the purine/pyrimidine and amino/keto encodings, repeat removal consistently reduced chromosome-level H, indicating that repetitive sequence substantially amplifies scaling in these representations. In contrast, the weak/strong encoding showed little net reduction after repeat exclusion, suggesting that repeat architecture is not a dominant driver of scaling under this encoding. Across all encodings, however, repeat-excluded chromosomes and low- or medium-repeat windows generally retained H values above 0.5, indicating that persistent correlation also extends beyond the most repeat-rich compartments.

### Shuffle-surrogate analysis confirms that LRCs exceed composition-only expectations

To test whether the observed LRD could be explained by nucleotide composition, we compared the wavelet-estimated Hurst exponent for each chromosome against complete shuffle-surrogate sequences that preserve purine/pyrimidine counts while destroying positional organization. Statistical significance was assessed using one-sided empirical surrogate tests, with BH-FDR correction as described in Methods.

Across chromosomes, we observed that H values are consistently higher than the shuffle-surrogate distributions and remained above the uncorrelated reference value of H = 0.5 (**Figure 5A**). The difference between observed and mean surrogate H was positive for all analysed chromosomes, with a mean ΔH of 0.284 across chromosomes. All 20 analysed chromosomes were significant after Benjamini–Hochberg correction (p^emp^=0.001, q^FDR^=0.001). Because p=0.001 is the minimum attainable empirical p-value with 999 surrogates, this indicates that no complete-shuffle surrogate reached or exceeded the observed chromosome-level H. These results show that the observed correlation signal cannot be attributed solely to global nucleotide composition or finite sequence length, and instead reflects sequence ordering beyond composition alone.

**Figure 5:**
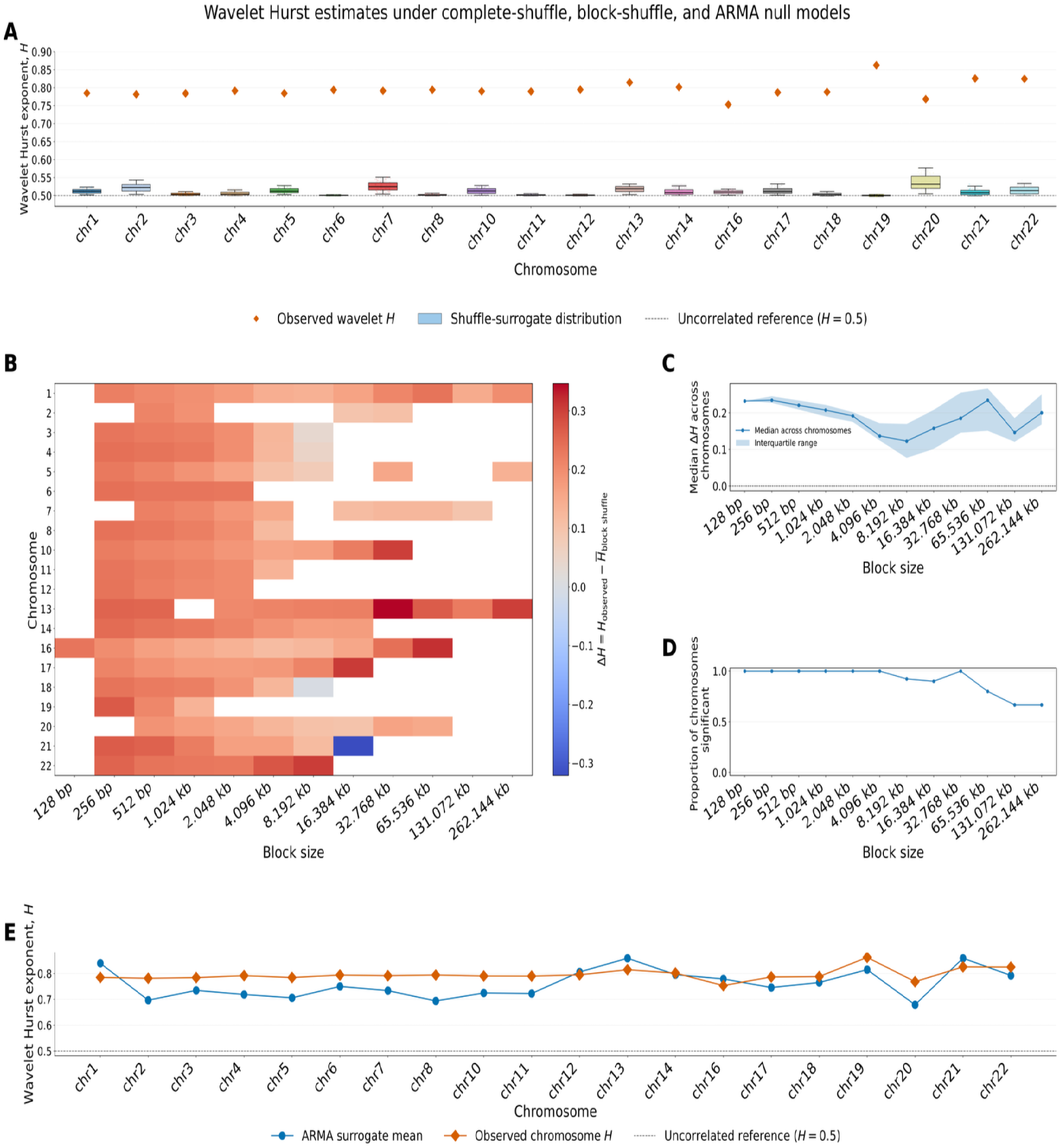
Surrogate-based assessment of chromosome-level LRDs. **(A)** Shuffle-surrogate distributions of wavelet Hurst exponent for each chromosome, with observed values as diamonds; **(B)** Observed H minus block-shuffle mean H; (**C)** Across-chromosome block-shuffle effect size; **(D)** Proportion of chromosomes whose observed H exceeded the 97.5th percentile of the corresponding block-shuffle surrogate distribution; **(E)** ARMA short-memory surrogate analysis. Observed chromosome-level wavelet Hurst exponents were compared with surrogate distributions generated from chromosome-specific BIC-selected ARMA models.

The complete-shuffle analysis was also repeated using the amino/keto and weak/strong encodings. Under the amino/keto encoding, observed chromosome-level wavelet H values were significantly greater than the corresponding complete-shuffle expectations for 21 of the 22 monofractal chromosomes after BH-FDR correction. For these 21 chromosomes, we found p^emp^=0.001 and q^FDR^=0.001. Under the weak/strong encoding, all 15 analysed monofractal chromosomes were significant after BH-FDR correction, with p^emp^=0.001 and q^FDR^=0.001. These results indicate that the excess correlation relative to composition-preserving shuffle nulls is not specific to the primary purine/pyrimidine encoding.

### Block-shuffle surrogates show that LRCs are not explained by local sequence structure alone

We next evaluated whether the observed chromosome-level scaling could be reproduced after preserving local sequence structure within genomic blocks. For each valid chromosome and block size, we compared the observed wavelet Hurst exponent with the corresponding block-shuffle surrogate distribution using a one-sided empirical surrogate test, followed by Benjamini–Hochberg correction.

The resulting ΔH was generally positive across chromosomes and retained block sizes (**Figure 5B**), indicating that the observed H values were typically higher than expected when only within-block structure was preserved. Overall, 129 of 133 chromosome–block-size combinations were significant after global BH-FDR correction. The median ΔH was positive at every retained block size, ranging from approximately 0.123 to 0.235, with 10 of 12 block sizes showing positive ΔH for all tested chromosomes. This supports the conclusion that the observed LRC signal is not explained solely by local composition, clustering, or short-range structure within genomic blocks. The across-chromosome median ΔH remained positive across the tested block sizes, although the effect size varied with block length (**Figure 5C**). This pattern is consistent with the block-shuffle null becoming more stringent as larger blocks preserve increasingly extended segments of the original sequence.

To test whether the block-shuffle effect was consistently positive across chromosomes, we applied a one-sided Wilcoxon signed-rank test to ΔH at each block size, followed by Benjamini–Hochberg correction across block sizes. The across-chromosome positive shift was significant for 8 of 12 retained block sizes after FDR correction, including block sizes from 256 bp to 8,192 bp, 32,768 bp, and 65,536 bp. The non-significant block sizes were either represented by very few valid chromosomes, such as 128 bp, 131,072 bp, and 262,144 bp, or showed one chromosome-specific deviation at 16,384 bp. Thus, the loss of significance at some larger block sizes should be interpreted in the context of reduced chromosome counts and more stringent local preservation, rather than as a general disappearance of the effect.

As a conservative descriptive summary, **Figure 5D** shows the proportion of chromosomes whose observed Hurst exponent exceeded the 97.5th percentile of the corresponding block-shuffle surrogate distribution. A substantial proportion of chromosomes remained above this upper-tail threshold across retained block sizes. Together, these results show that the observed LRC signal is not explained solely by local composition, clustering or short-range structure preserved within genomic blocks. Instead, the persistence of elevated H values under block-shuffle randomization supports the presence of sequence ordering beyond the local block scale.

Chr21 was the main exception to the otherwise positive block-shuffle pattern. In the heatmap, chr21 showed a negative ΔH at the largest retained block size (**Figure 5B**). The chr21 diagnostic assessment showed that this deviation was restricted to this block-size condition: observed H exceeded the block-shuffle mean at smaller and intermediate block sizes, but fell below it at L=16 kb (**Supplementary Figure 5**). This same condition corresponded to a narrow adjusted fitting interval and perfect wavelet-fit (R^2^=1) values, indicating limited scale support for the slope estimate. Therefore, the chr21 result needs to be interpreted as a scale-interval-specific deviation rather than as a consistent chromosome-wide reversal of the block-shuffle pattern.

The block-shuffle analysis was also repeated using the complementary weak/strong and amino/keto encodings. Under the weak/strong encoding, 123 of 124 valid chromosome–block-size combinations remained significant after global BH-FDR correction, with positive median ΔH at all 10 retained block sizes, ranging from 0.153 to 0.267. The only exception was chrY at a block size of 1,024 bp, for which the observed H was lower than the block-shuffle mean. Under the amino/keto encoding, ΔH was positive in all 117 valid comparisons, and 114 remained significant after global BH-FDR correction. Median ΔH was positive across all 12 retained block sizes, ranging from 0.035 to 0.315, although the across-chromosome Wilcoxon test remained significant after FDR correction for 6 of the 12 block sizes, with reduced support at larger block sizes where fewer chromosomes retained valid fitting intervals. Together, these complementary analyses support the conclusion that the observed scaling generally extends beyond the locally preserved block scale across nucleotide encodings, while also showing that the strength and consistency of this effect depend on the encoding and available scale support.

### ARMA short-memory surrogates partially reproduce chromosome-level H but do not uniformly explain the observed scaling

To evaluate whether elevated chromosome-level Hurst exponents could be reproduced by conventional short-memory autocorrelation, we compared the observed wavelet H values with 999 ARMA surrogates generated from chromosome-specific BIC-selected ARMA models. For the purine/pyrimidine encoding, ARMA analyses were restricted to the 20 monofractal chromosomes, excluding chr9, chr15, chrX, and chrY, which were analysed separately using MF-DFA.

ARMA surrogates produced elevated H values across many chromosomes (**Figure 5E**), unlike complete-shuffle surrogates, which remained close to the uncorrelated expectation of H = 0.5. The observed H exceeded the ARMA surrogate mean in 15 of 20 chromosomes and exceeded the ARMA 97.5th percentile in the same 15 chromosomes. These 15 chromosomes were significant after Benjamini-Hochberg correction, with empirical p-values of 0.001-0.002 and FDR-adjusted q-values of 0.0014-0.0027. Across all chromosomes, the difference between observed and mean ARMA-surrogate H ranged from -0.055 to 0.100, with a median ΔH of 0.046. However, the ARMA null did not provide a uniform explanation across chromosomes. In five chromosomes, chr1, chr12, chr13, chr16, and chr21, the ARMA surrogate mean was comparable to or higher than the observed H. This indicates that finite-order short-memory autocorrelation can reproduce or exceed the observed H in some chromosome-specific scale intervals. Therefore, ARMA surrogates provide a conservative short-memory comparison rather than a complete genome-wide rejection of the short-memory null.

Repeating the ARMA analysis with the complementary encodings produced similarly heterogeneous, but encoding-dependent, results (**Supplementary Figures 8-9**). Under the amino/keto encoding, the observed H exceeded the ARMA surrogate mean in 12 of 22 monofractal chromosomes and exceeded the surrogate 97.5th percentile in 11, with ΔH ranging from −0.097 to 0.304 and a median of 0.029. Under the weak/strong encoding, the observed H exceeded both the surrogate mean and the 97.5th percentile in only 4 of 15 monofractal chromosomes, chr5, chr6, chr8, and chr19, while the ARMA mean was equal to or greater than the observed value in the remaining 11 chromosomes; median ΔH was −0.007. These findings reinforce that finite-order short-memory models can reproduce elevated H over many chromosome-specific scale intervals, although their explanatory capacity varies substantially with both chromosome and nucleotide encoding.

Together with the complete-shuffle and block-shuffle analyses, these results support a nuanced interpretation: the observed correlation structure cannot be attributed to composition alone or local sequence clustering alone, but short-memory autocorrelation may contribute to elevated H in some chromosomes. Thus, the full pattern supports a heterogeneous mixture of long-range organization, local structure, repeat architecture, and chromosome-specific short-memory effects.

## Discussion

Our analysis revisits LRCs in the human genome using the first complete T2T reference genome, allowing chromosome-scale correlation structure to be evaluated across regions that were previously missing or fragmented. We find that most chromosomes exhibit persistent LRD over chromosome-specific scaling intervals, supporting earlier observations that genomic DNA contains statistical dependencies extending beyond local sequence composition (Peng et al. 1992, 1994; Arneodo et al. 1995). However, the complete T2T assembly reveals that this organization is not uniform across chromosomes. Instead, the strength, scale range, and complexity of LRCs vary substantially, consistent with a genome organized as a mosaic of compositional domains, repeat-rich compartments, and structurally distinct chromosome regions.

A central implication of our results is that chromosome-level Hurst exponents should not be interpreted as uniform properties of entire chromosomes. Although most chromosomes showed persistent scaling over chromosome-specific intervals, the sliding-window Hurst landscapes and genome-wide scalograms demonstrated that this signal is heterogeneous and multifactorial. Thus, a single chromosome-wide exponent should be viewed as a summary statistic of a range of Hurst exponents rather than as evidence for a single homogeneous correlation. This interpretation is consistent with previous work arguing that LRCs in DNA may arise from a mixture of compositional domains, including patchiness, repeats, and multiple characteristic length scales rather than from one process (Karlin and Brendel 1993; Larhammar and Chatzidimitriou-Dreismann 1993; Carpena et al. 2007). In our work, local Hurst exponent values differed across CenSat annotation classes, whereas gene-feature categories showed weak effects. This suggests that repeat architecture and centromere-associated sequence organization explain a larger fraction of the strongest local variation in scaling behavior than genic content. These observations are also consistent with previous work linking repeats and large-scale genomic organization to correlation structure in human DNA (Holste et al. 2003).

For the purine/pyrimidine encoding, chromosomes 9, 15, X, and Y were notable because their fluctuation profiles were not adequately captured by a single monofractal scaling regime. Instead, ML-DFA selected quartic or otherwise complex profiles, and MF-DFA revealed strong q-dependence of the generalized Hurst exponent, nonlinear mass exponents, and broad singularity spectra. Similar chromosome-specific results were also observed for the weak/strong and amino/keto encodings. These results indicate that chromosomes contain heterogeneous scaling regimes in which small- and large-fluctuation regions behave differently across genomic scales.

However, the repeat-exclusion analysis also shows that repeats do not fully explain the signal. After annotated repeats were removed, chromosome-level H remained above the uncorrelated expectation of H = 0.5 for all analysed chromosomes. Therefore, the observed LRC cannot be attributed exclusively to repeat sequences. Instead, the results support a layered model in which different genomic compartments contribute at different scales: satellite-rich and repeat-dense regions amplify local and chromosome-level persistence, while non-repeat sequence retains additional correlation structure, possibly reflecting broader compositional domains, chromosome-arm organization, isochores, duplicated sequence, or other forms of regional sequence organization.

This distinction is important for interpreting LRCs in the T2T genome. Earlier incomplete assemblies limited access to the most repeat-rich and satellite-rich regions, whereas the T2T assembly allows these compartments to be included directly. Our results suggest that the complete representation of repetitive DNA does not simply add noise to LRC analysis; rather, it reveals that repeats are an integral component of chromosome-scale sequence organization. At the same time, because repeat-excluded chromosomes retain elevated H, the genome-wide LRC signal should not be reduced to a repeat-only phenomenon. A more accurate interpretation is that LRC in human chromosomes emerges from the combined organization of repetitive, satellite-rich, and non-repeat sequence compartments.

Surrogate null models provide an additional layer of interpretation by asking whether elevated chromosome-level Hurst exponents could be reproduced by simpler non-LRD mechanisms. Complete-shuffle surrogates preserved the global purine/pyrimidine composition but destroyed positional order, and therefore tested whether unequal nucleotide-class frequencies alone were sufficient to explain the observed scaling. The fact that observed H values exceeded the shuffle-surrogate distributions across chromosomes indicates that composition alone is not sufficient. Block-shuffle surrogates provided a stronger local-structure null by preserving sequence organization within genomic blocks while disrupting the ordering of those blocks along chromosomes. The persistence of elevated observed H values relative to block-shuffle expectations indicates that local clustering and within-block structure alone do not fully account for the chromosome-level signal.

The ARMA surrogate analysis further refines this interpretation by testing a conventional short-memory alternative. Stationary ARMA processes can exhibit autocorrelation, but their autocorrelation decays exponentially rather than as the slower power-law decay expected under LRD. Therefore, ARMA surrogates ask whether finite-order short-memory autocorrelation can mimic the observed wavelet H values over the finite scale intervals analysed here. In contrast to the complete-shuffle and block-shuffle results, the ARMA null produced a more mixed pattern: ARMA surrogates generated elevated H values for several chromosomes, and the observed H did not uniformly exceed the ARMA surrogate distribution across all chromosomes. This result does not invalidate the evidence for chromosome-scale correlation structure, but it shows that some apparent persistence can be reproduced by short-memory models over finite genomic ranges. Thus, the ARMA analysis supports a cautious interpretation: elevated H should not be treated as definitive proof of asymptotic LRD for every chromosome, but as part of a broader evidence chain that also includes scale segmentation, spatial H landscapes, repeat-exclusion analysis, multifractal behaviour, and composition/local-structure-preserving null models.

Several limitations should be considered. First, the analysis is based on a single reference genome, and therefore does not yet capture population-level variation in LRC structure. Extending this framework to complete diploid assemblies and pangenome resources will be important for determining whether LRC landscapes vary among individuals, ancestries, haplotypes, or structurally polymorphic regions (Nurk et al. 2022). Second, the purine/pyrimidine encoding captures one biologically meaningful encoding but other encodings may have different autocorrelation patterns. Third, Hurst exponents estimated from finite chromosomes should be interpreted as scale-specific summaries rather than direct proof of asymptotic LRD. This is particularly important because short-memory processes, compositional heterogeneity, and finite-size effects can produce elevated H estimates over restricted scale intervals. Fourth, repeat exclusion is a useful sensitivity analysis but does not reconstruct a biologically realistic repeat-free chromosome, because removing annotated repeats changes genomic spacing and creates artificial junctions between formerly distant non-repeat regions.

In summary, the complete T2T human genome reveals that LRC is a widespread but spatially heterogeneous property of human chromosomes and encodings. Rather than supporting a single uniform genome-wide fractal model, our results suggest that the human genome is organized as a mosaic of correlation regimes shaped by multiple encodings, inter- and intra-chromosomal differences, and long-range sub-compartmentalizations. Repeat exclusion shows that annotated repeats substantially elevate chromosome-level H, but the persistence of H > 0.5 after repeat removal indicates that non-repeat sequence also contributes to long-range organization. Surrogate analyses further show that this signal is not explained by composition alone or by local sequence structure alone, while ARMA surrogates indicate that finite short-memory autocorrelation can account for part of the observed H range in some chromosomes. Thus, our interpretation is that LRCs in the T2T genome arise from a heterogeneous combination of repeat architecture, satellite-rich sequence, broader compositional domains, and chromosome-specific sequence organization, rather than from a single universal mechanism. This framework provides a basis for future studies linking long-range sequence organization to genome evolution, structural variation, recombination, chromatin architecture, and disease-associated instability.

## Data and Code availability

All scripts developed for this study can be found on the following GitHub repository: https://github.com/Georgakopoulos-Soares-lab/dna_long_range_correlations

The T2T-CHM13v2.0 human reference genome assembly analysed in this study is publicly available through the NCBI Assembly database under accession **GCF_009914755.1**. All genomic annotation datasets used in the analyses are publicly available from the Human Pangenome Reference Consortium T2T-CHM13 resource.

Cytoband annotations are available at:

https://s3-us-west-2.amazonaws.com/human-pangenomics/T2T/CHM13/assemblies/annotation/chm13v2.0_cytobands_allchrs.bed

GENCODE v35 CAT/Liftoff gene annotations are available at:

https://s3-us-west-2.amazonaws.com/human-pangenomics/T2T/CHM13/assemblies/annotation/chm13v2.0_GENCODEv35_CAT_Liftoff.vep.gff3.gz

Centromeric and satellite annotations from CenSat v2.1 are available at:

https://s3-us-west-2.amazonaws.com/human-pangenomics/T2T/CHM13/assemblies/annotation/chm13v2.0_censat_v2.1.bed

RepeatMasker annotations are available at:

https://s3-us-west-2.amazonaws.com/human-pangenomics/T2T/CHM13/assemblies/annotation/chm13v2.0_RepeatMasker_4.1.2p1.2022Apr14.bed

## Contributions

K.P. developed the methodological framework, designed and implemented the computational analyses, curated and processed the data, performed the statistical analyses, interpreted the results, generated the figures and tables, and wrote the Methods and Results sections of the manuscript. I.G-S, conceived the study, wrote the initial drafts of the Abstract, Introduction, and Discussion and contributed to the interpretation and contextualization of the findings. K.P. and I.G.S. wrote the manuscript with help from G.M. and A.Z..

## Acknowledgments

Research reported in this publication was supported by the National Institute of General Medical Sciences under award number R35GM155468 (to I.G.S.)

## Supplementary Figures and Tables

**Supplementary Table 1:**
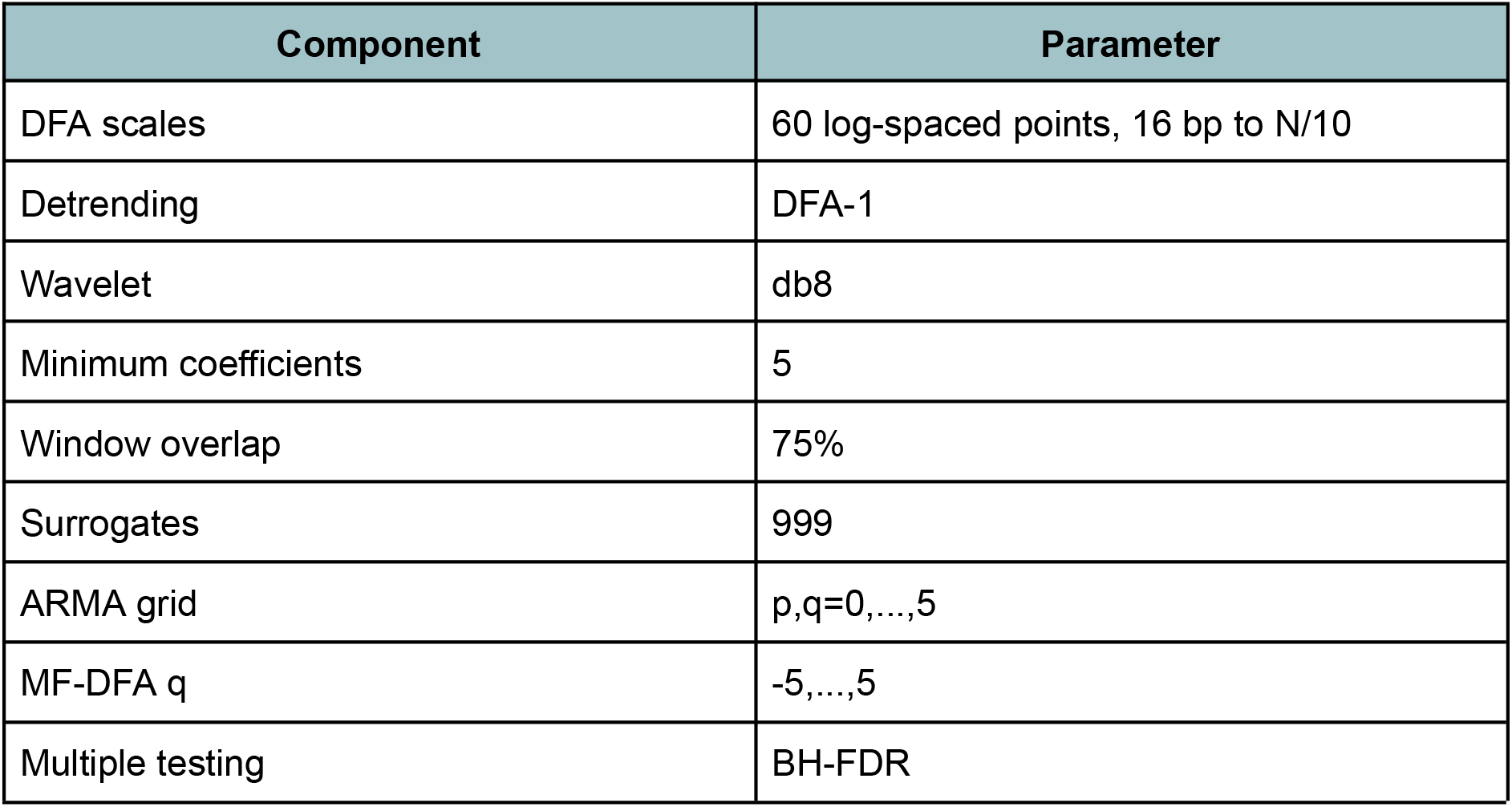
Summary of the principal analytical parameters used throughout the study. The table lists the scale ranges, detrending settings, wavelet configuration, sliding-window parameters, surrogate sample size, ARMA order-search grid, MF-DFA moment range, and multiple-testing correction applied in the chromosome-level and local long-range correlation analyses.

**Supplementary Figure 1:**
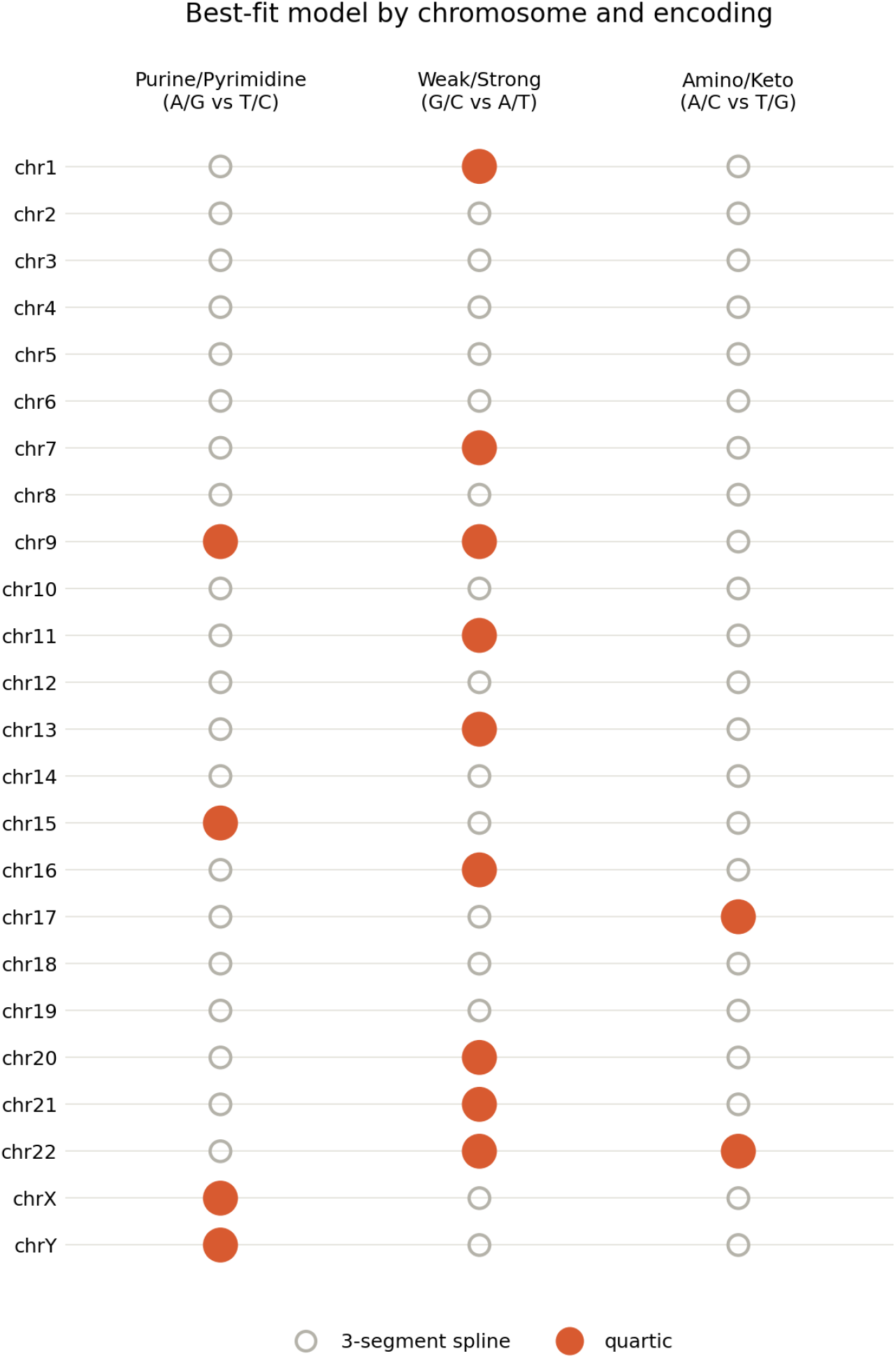
Best ML DFA fitted for each chromosome for the three different encodings.

**Supplementary Figure 2:**
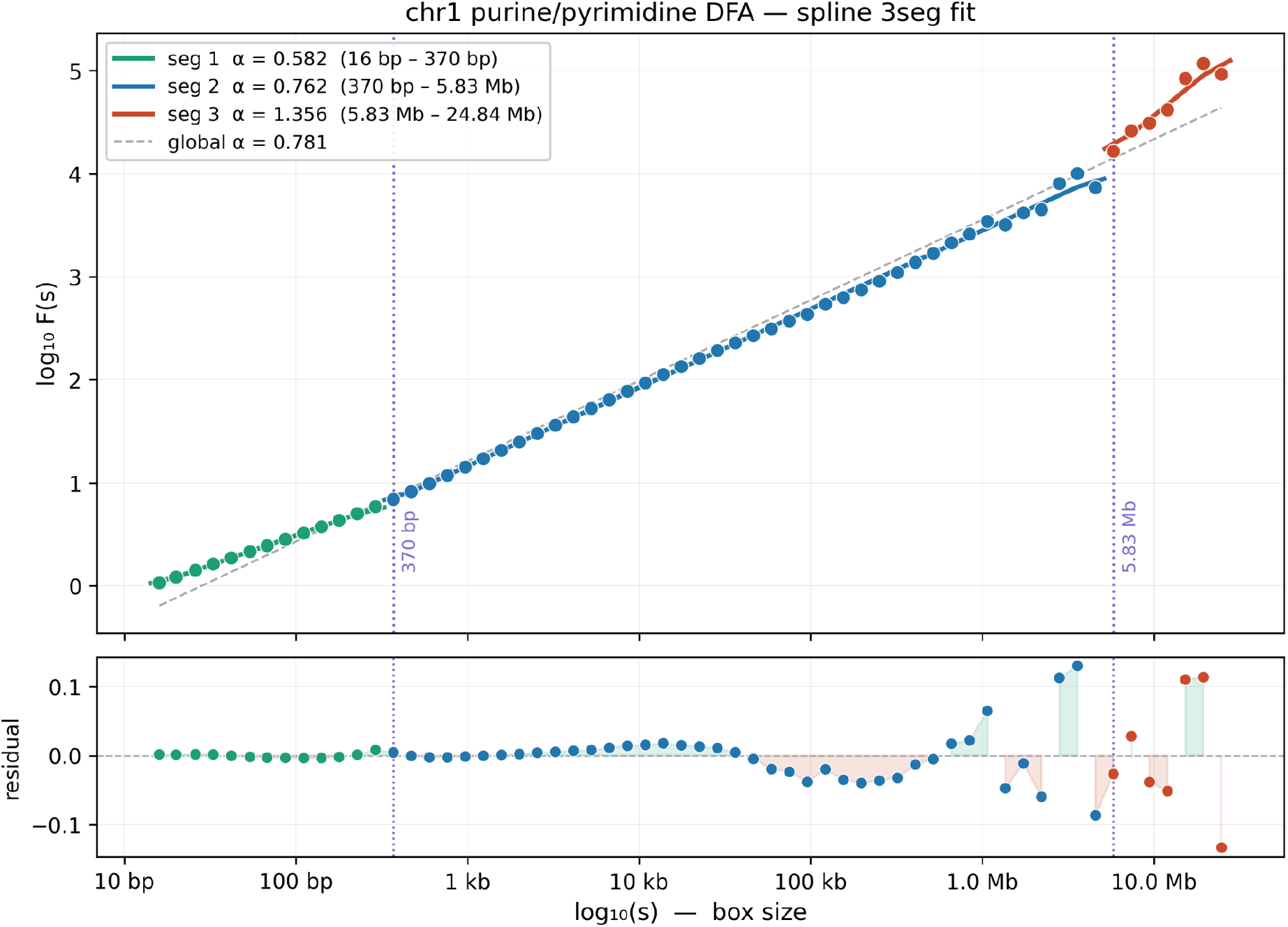
Three-segment spline fit of the ML-DFA fluctuation function for the purine/pyrimidine encoding of chromosome 1. The log-log relationship between fluctuation magnitude F(s) and box size identifies three scaling regimes with exponents α=0.582, 0.762, and 1.356, separated by breakpoints at 370 bp and 5.83 Mb. The lower panel shows the residuals of the piecewise fit, indicating a good overall fit across scales.

**Supplementary Figure 3:**
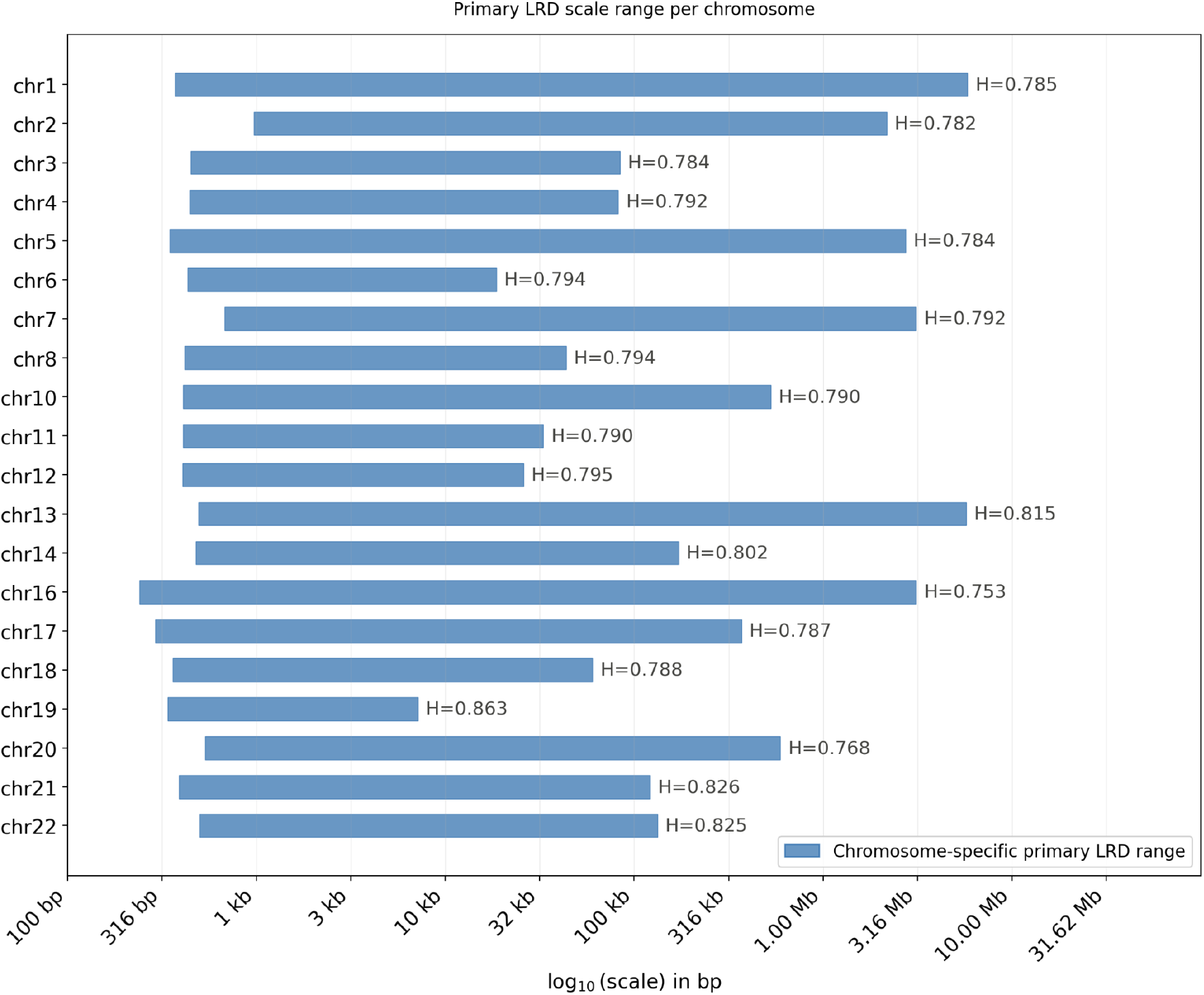
Chromosome-specific primary LRD scale intervals in the complete human T2T genome. For each chromosome, ML-DFA was used to find the scaling intervals supporting persistent long range correlations.

**Supplementary Figure 4:**
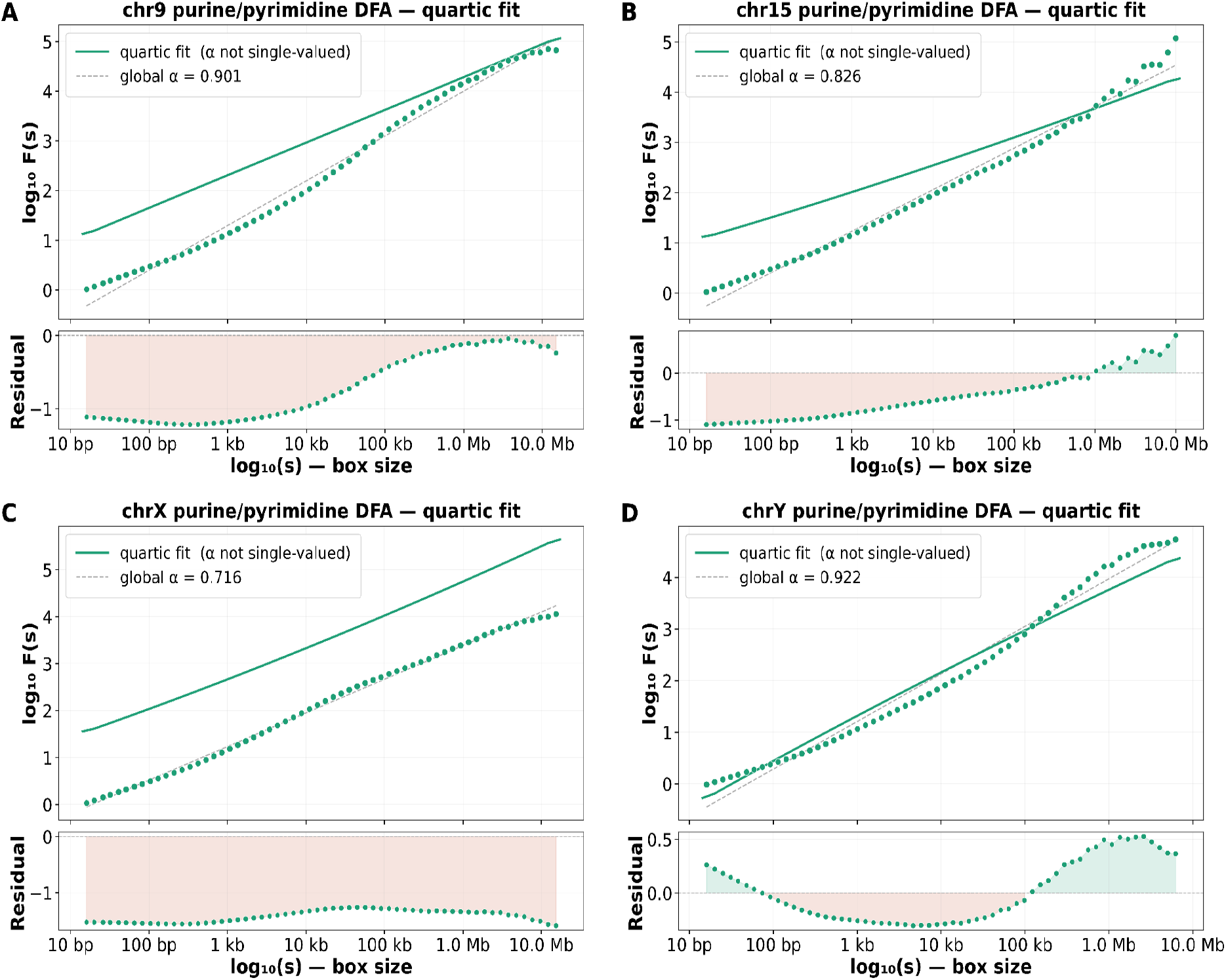
Chromosomes 9, 15, X, and Y were classified by ML-DFA as exhibiting quartic or otherwise complex scaling behaviour.

**Supplementary Figure 5:**
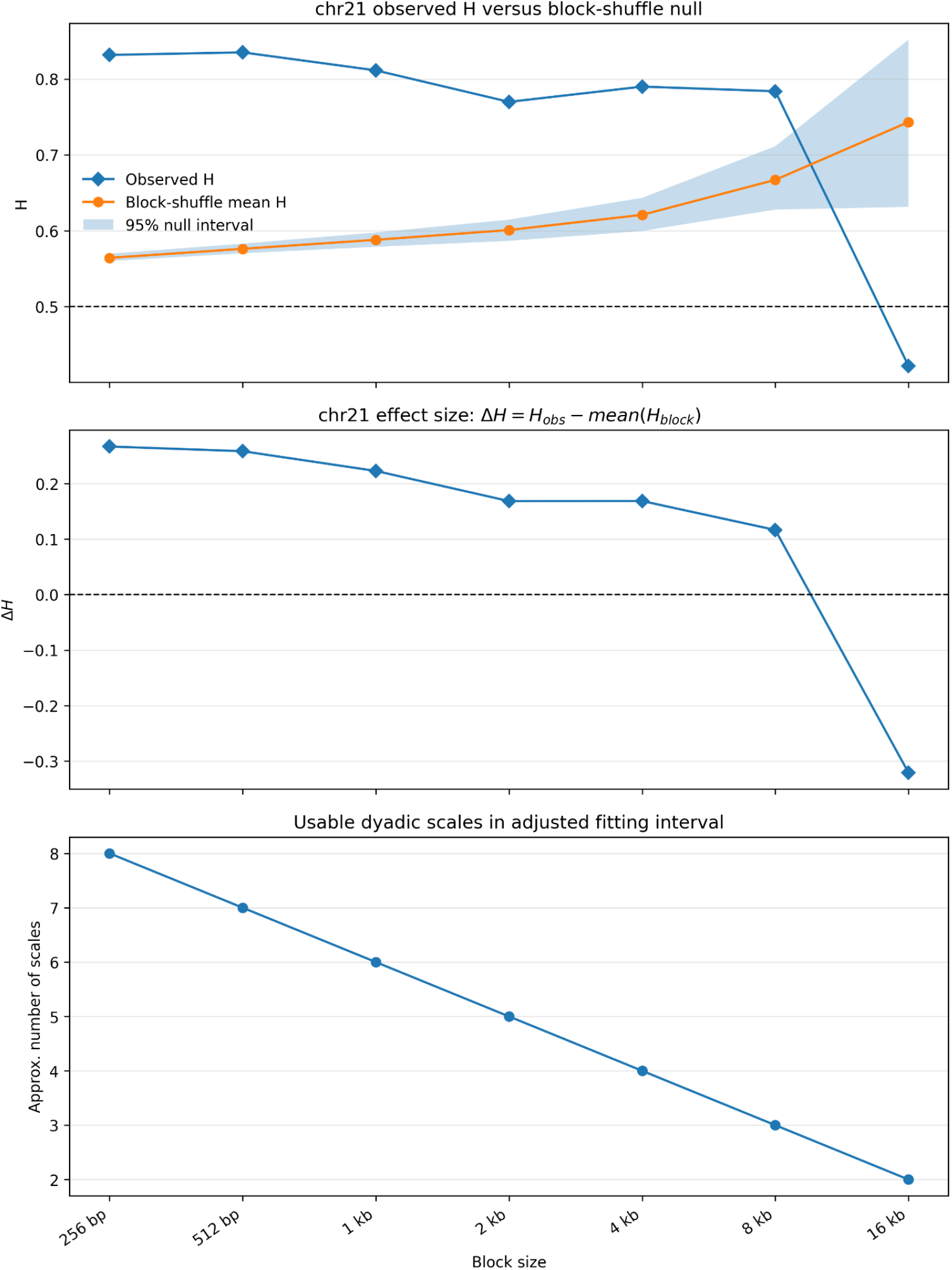
Block-shuffle sensitivity analysis for chromosome 21. Observed wavelet H, block-shuffle null mean and 95% interval, corresponding effect size ΔH, and the number of usable dyadic scales are shown across increasing block sizes. The observed signal exceeds the null up to 8 kb, while the 16 kb estimate is unreliable because only two fitting scales remain.

**Supplementary Figure 6:**
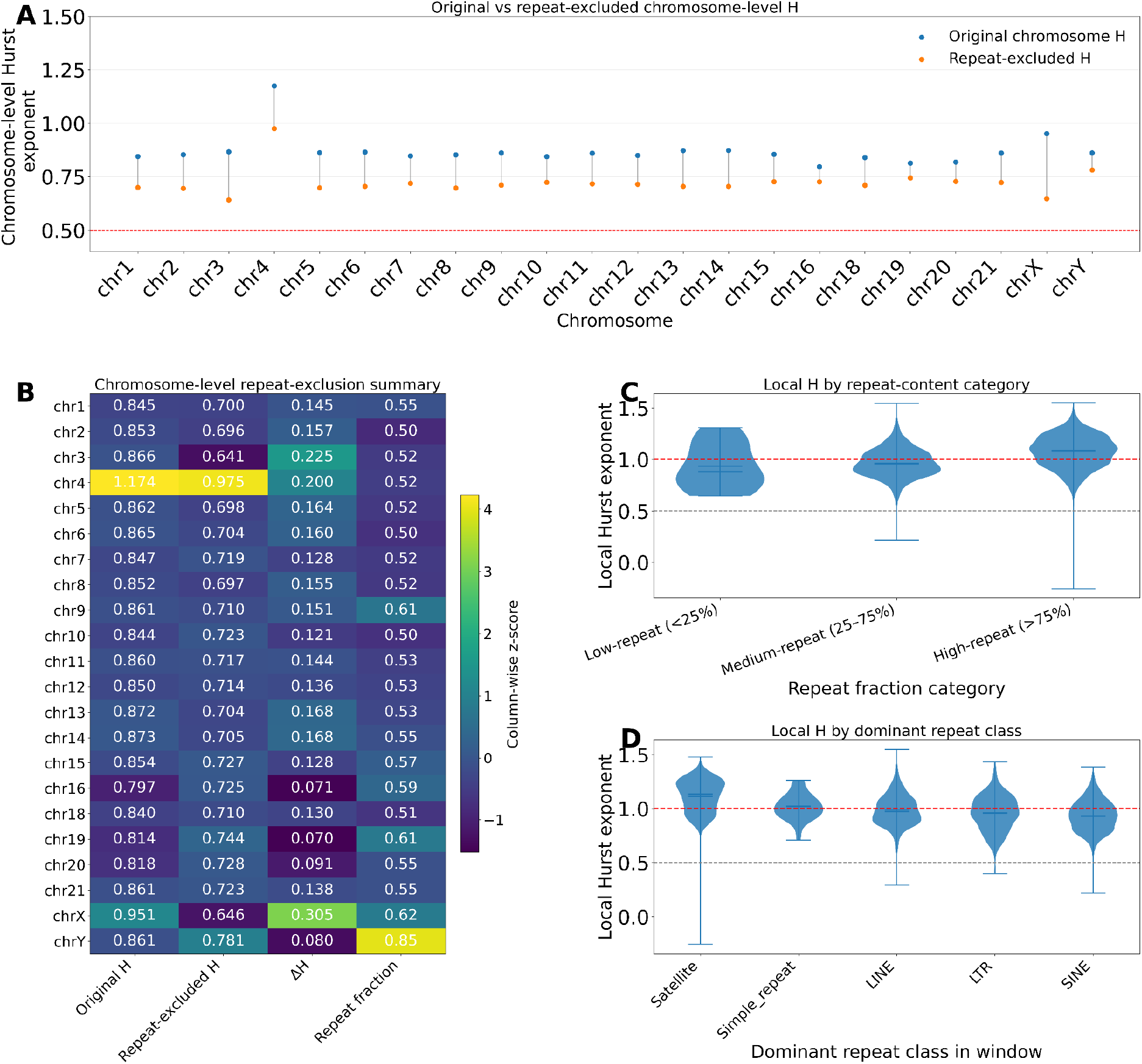
Effects of repeat content on chromosome-level and local Hurst exponents with the amino/keto encoding.

**Supplementary Figure 7:**
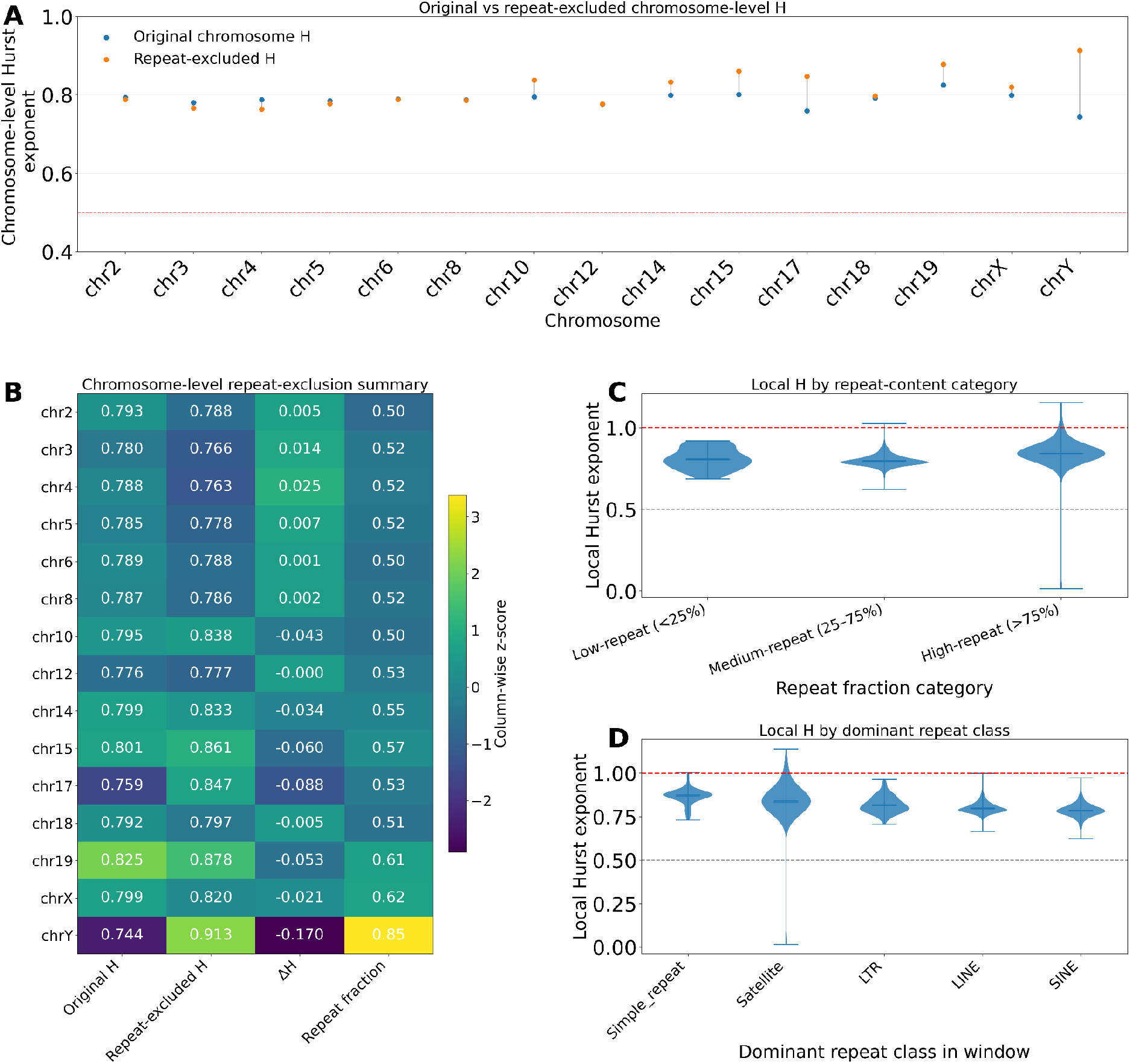
Effects of repeat content on chromosome-level and local Hurst exponents with the weak/strong encoding.

**Supplementary Table 2:** Summary results across all chromosomes for the amino/keto encoding. ML-DFA best model selection, scaling interval, DFA scaling exponent α, Wavelet Hurst exponent H and R^2^ goodness of fit are shown. A dash indicates that the corresponding monofractal estimate was not applicable.

| Chromosome | ML-DFA model | Scaling interval | $\alpha$ | $H_{\text{wav}}$ | $R^2$ |
| --- | --- | --- | --- | --- | --- |
| chr1 | Three-segment spline | 1,577 bp–9.45 Mb | 0.827 | 0.845 | 0.981 |
| chr2 | Three-segment spline | 1,566 bp–4.48 Mb | 0.824 | 0.853 | 0.983 |
| chr3 | Three-segment spline | 1,473 bp–84.28 kb | 0.842 | 0.866 | 0.980 |
| chr4 | Three-segment spline | 132.38 kb–19.36 Mb | 0.897 | 1.741 | 0.946 |
| chr5 | Three-segment spline | 1,127 bp–3.48 Mb | 0.812 | 0.862 | 0.980 |
| chr6 | Three-segment spline | 1,107 bp–1.29 Mb | 0.800 | 0.865 | 0.983 |
| chr7 | Three-segment spline | 1,370 bp–1.54 Mb | 0.814 | 0.847 | 0.987 |
| chr8 | Three-segment spline | 1,054 bp–896.84 kb | 0.796 | 0.852 | 0.982 |
| chr9 | Three-segment spline | 1,695 bp–4.69 Mb | 0.842 | 0.861 | 0.989 |
| chr10 | Three-segment spline | 1,295 bp–1.06 Mb | 0.809 | 0.844 | 0.988 |
| chr11 | Three-segment spline | 1,296 bp–1.69 Mb | 0.806 | 0.860 | 0.983 |
| chr12 | Three-segment spline | 1,291 bp–1.05 Mb | 0.801 | 0.850 | 0.985 |
| chr13 | Three-segment spline | 1,935 bp–5.72 Mb | 0.856 | 0.872 | 0.986 |
| chr14 | Three-segment spline | 1,481 bp–5.13 Mb | 0.842 | 0.873 | 0.984 |
| chr15 | Three-segment spline | 1,848 bp–5.06 Mb | 0.846 | 0.854 | 0.988 |
| chr16 | Three-segment spline | 1,825 bp–409.57 kb | 0.796 | 0.797 | 0.985 |
| chr17 | Quartic | — | — | — | — |
| chr18 | Three-segment spline | 1,097 bp–557.59 kb | 0.785 | 0.840 | 0.983 |
| chr19 | Three-segment spline | 1,937 bp–3.21 Mb | 0.793 | 0.814 | 0.976 |
| chr20 | Three-segment spline | 1,597 bp–593.92 kb | 0.793 | 0.818 | 0.991 |
| chr21 | Three-segment spline | 1,126 bp–185.57 kb | 0.818 | 0.861 | 0.998 |
| chr22 | Quartic | — | — | — | — |
| chrX | Three-segment spline | 1,353 bp–28.16 kb | 0.934 | 0.951 | 0.988 |
| chrY | Three-segment spline | 1,258 bp–1.09 Mb | 0.825 | 0.861 | 0.972 |

**Supplementary Table 3:** Summary results across all chromosomes for the weak/strong encoding. ML-DFA best model selection, scaling interval, DFA scaling exponent α, Wavelet Hurst exponent H and R^2^ goodness of fit are shown. A dash indicates that the corresponding monofractal estimate was not applicable.

| Chromosome | ML-DFA model | Scaling interval | $\alpha$ | $H_{\text{wav}}$ | $R^2$ |
| --- | --- | --- | --- | --- | --- |
| chr1 | Quartic | — | — | — | — |
| chr2 | Three-segment spline | 110 bp–194.91 kb | 0.785 | 0.793 | 0.996 |
| chr3 | Three-segment spline | 107 bp–172.13 kb | 0.767 | 0.780 | 0.993 |
| chr4 | Three-segment spline | 107 bp–550.04 kb | 0.764 | 0.788 | 0.991 |
| chr5 | Three-segment spline | 106 bp–414.82 kb | 0.778 | 0.785 | 0.991 |
| chr6 | Three-segment spline | 83 bp–314.70 kb | 0.760 | 0.789 | 0.994 |
| chr7 | Quartic | — | — | — | — |
| chr8 | Three-segment spline | 103 bp–280.25 kb | 0.775 | 0.787 | 0.995 |
| chr9 | Quartic | — | — | — | — |
| chr10 | Three-segment spline | 128 bp–166.48 kb | 0.789 | 0.795 | 0.994 |
| chr11 | Quartic | — | — | — | — |
| chr12 | Three-segment spline | 102 bp–131.18 kb | 0.763 | 0.776 | 0.993 |
| chr13 | Quartic | — | — | — | — |
| chr14 | Three-segment spline | 154 bp–87.15 kb | 0.799 | 0.799 | 0.990 |
| chr15 | Three-segment spline | 193 bp–108.29 kb | 0.817 | 0.801 | 0.989 |
| chr16 | Quartic | — | — | — | — |
| chr17 | Three-segment spline | 187 bp–31.72 kb | 0.789 | 0.759 | 0.974 |
| chr18 | Three-segment spline | 95 bp–146.71 kb | 0.765 | 0.792 | 0.996 |
| chr19 | Three-segment spline | 651 bp–291.60 kb | 0.833 | 0.825 | 0.995 |
| chr20 | Quartic | — | — | — | — |
| chr21 | Quartic | — | — | — | — |
| chr22 | Quartic | — | — | — | — |
| chrX | Three-segment spline | 131 bp–367.65 kb | 0.776 | 0.799 | 0.979 |
| chrY | Three-segment spline | 525 bp–152.93 kb | 0.743 | 0.744 | 0.931 |

**Supplementary Figure 8:**
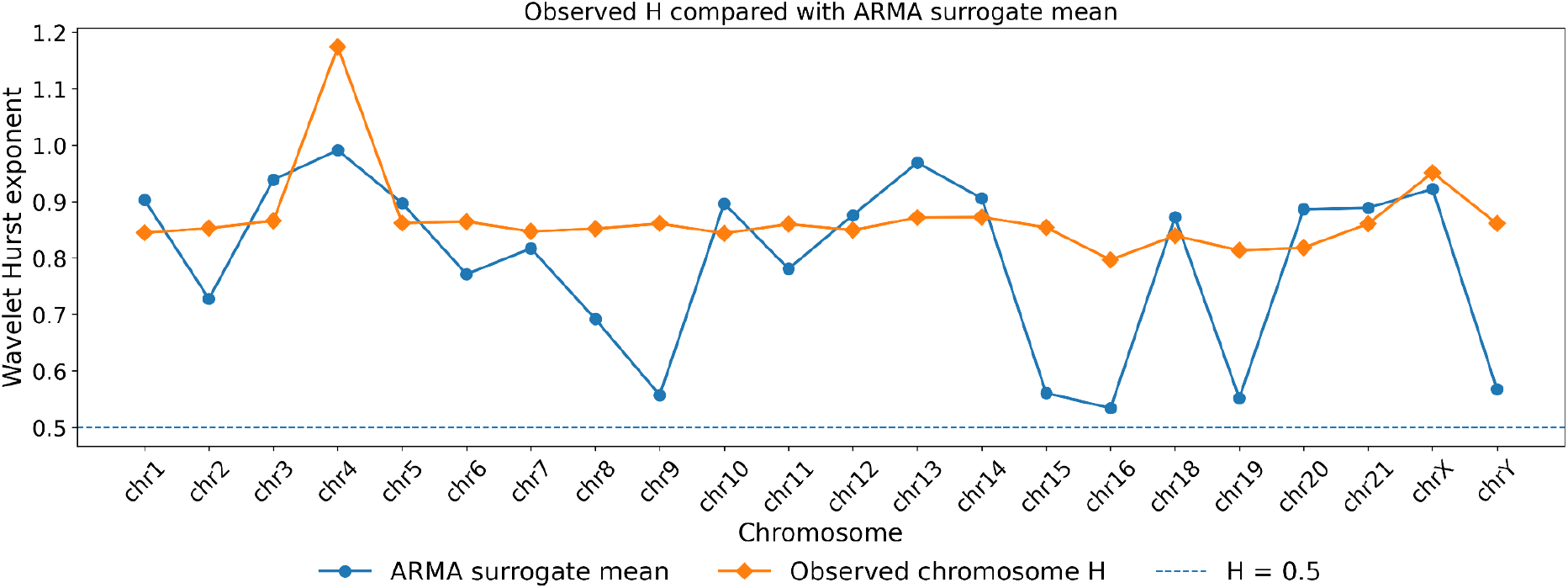
ARMA short-memory surrogate analysis for the **amino/keto** encoding. Observed chromosome-level wavelet Hurst exponents were compared with surrogate distributions generated from chromosome-specific BIC-selected ARMA models.

**Supplementary Figure 9:**
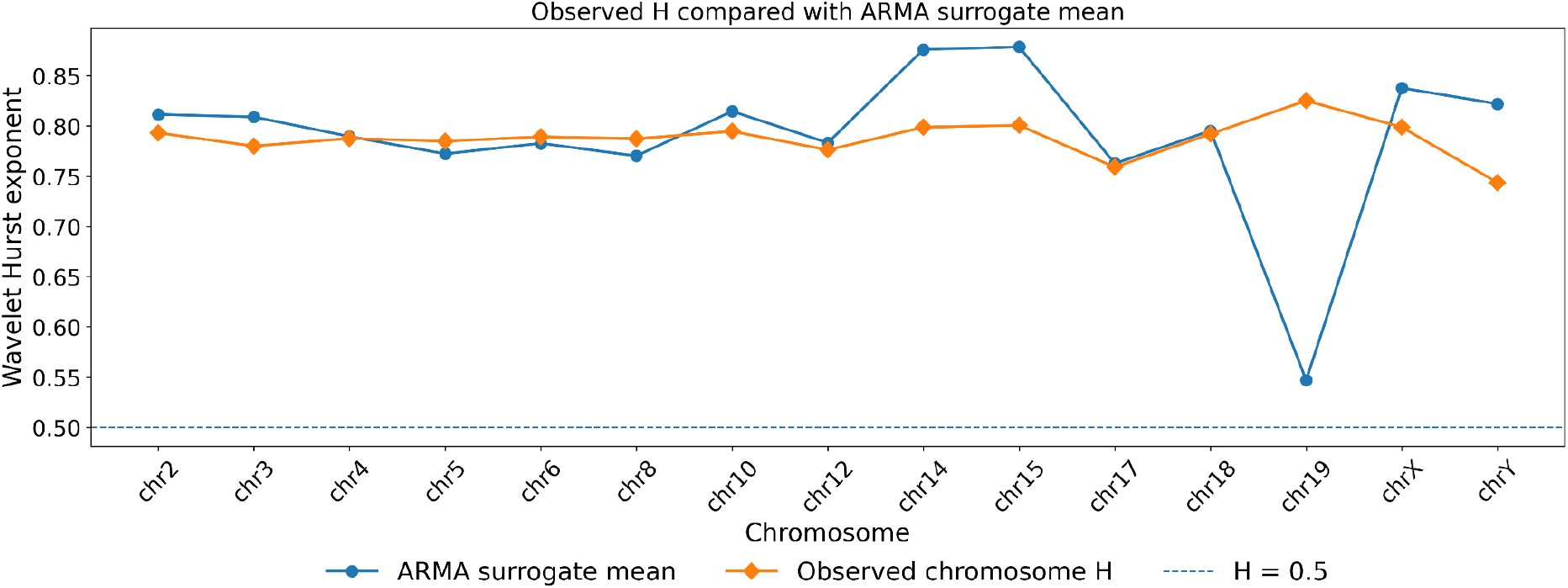
ARMA short-memory surrogate analysis for the **weak/strong** encoding. Observed chromosome-level wavelet Hurst exponents were compared with surrogate distributions generated from chromosome-specific BIC-selected ARMA models.

